# An internal expectation guides *Drosophila* egg-laying decisions

**DOI:** 10.1101/2021.09.30.462671

**Authors:** Vikram Vijayan, Zikun Wang, Vikram Chandra, Arun Chakravorty, Rufei Li, Stephanie L. Sarbanes, Hessameddin Akhlaghpour, Gaby Maimon

## Abstract

When presented with two egg-laying substrates, *Drosophila* lay most of their eggs on the option with higher relative value. How do flies make this relative-value decision? Might the strategy they use allow them to choose the best option even when they experience substrates with a more complex spatiotemporal profile than in canonical two-choice paradigms? We measured *Drosophila* egg-laying behavior in diverse substrate environments. In all cases, we found that flies dynamically increase or decrease their egg-laying rates as they explore substrates for a deposition site so as to target eggs to the best, recently visited option. Visiting the best option typically led to a strong reduction in egg laying on other substrates for several minutes, with this timescale varying across fly strains. Our data support a model in which flies compare the value of the current substrate with an internally constructed *expectation* on the value of available options to regulate the likelihood of laying an egg. We show that dopamine-neuron activity is critical for learning and/or expressing this expectation, similar to its role in certain tasks in vertebrates. Integrating sensory experiences over minutes to generate an internal sense of the quality of available options, i.e., forming an expectation, allows flies to use a dynamic reference point for judging the current substrate and might be a general way in which decisions are made, even beyond flies and egg laying.

## Introduction

When trying to identify the best fruit in a basket, each inspected item updates our internal expectation of what the basket has to offer. We use this expectation to settle for an unripe option, for example, if none of the items inspected over the past few seconds to minutes have been ripe. The process of updating one’s expectations based on recent experiences is central to many daily decisions. Might we be able to study such an expectation process in a tractable model system?

Female *Drosophila* lay dozens of eggs, one at a time, over a few hours. During the past decades, researchers have studied *Drosophila*’s egg laying preferences across a broad array of conditions^1–12^. In foundational work on sucrose substrates, it was shown that flies do not simply express innate preferences for specific sucrose concentrations. Instead, flies treat the same sucrose substrate as attractive or repulsive depending on the nature of a second nearby option^5,6^. This observation suggests that flies can assess the *relative value* of two sucrose substrates in deciding where to lay eggs. To better understand this relative-valuation process, we performed a detailed analysis of the egg laying behavior of *Drosophila* in simple two-choice chambers as well as in chambers with more than two substrate options or where they experienced options for very different lengths of time. The results argue that *Drosophila* form an internal expectation about their environment––akin to the expectation formed by humans in the fruit basket example––which guides their egg-laying decisions. The impact of this expectation on behavior is evident, typically, for few minutes after flies encounter a new substrate. Behavioral-genetic experiments implicate dopamine neurons in the expectation process.

## Results

### Flies can make a relative-value decision between two sucrose-containing substrates

We placed single wild type Canton-S (CS) flies, overnight, in dark, custom egg-laying substrate-choice chambers where agarose-based substrate islands are separated by a 2.5 mm wide plastic barrier^13^ (Fig. 1a, Fig. S1a). The 2.5 mm barrier is approximately the length of the fly, which means that the fly’s legs touched either one substrate or the other, but rarely (if ever) touched both substrates at the same time. In all experiments, agarose substrates contained 1.6% ethanol and 0.8% acetic acid, simulating a rotting fruit^1^. Substrates additionally contained varying amounts of sucrose.

**Fig. 1.**
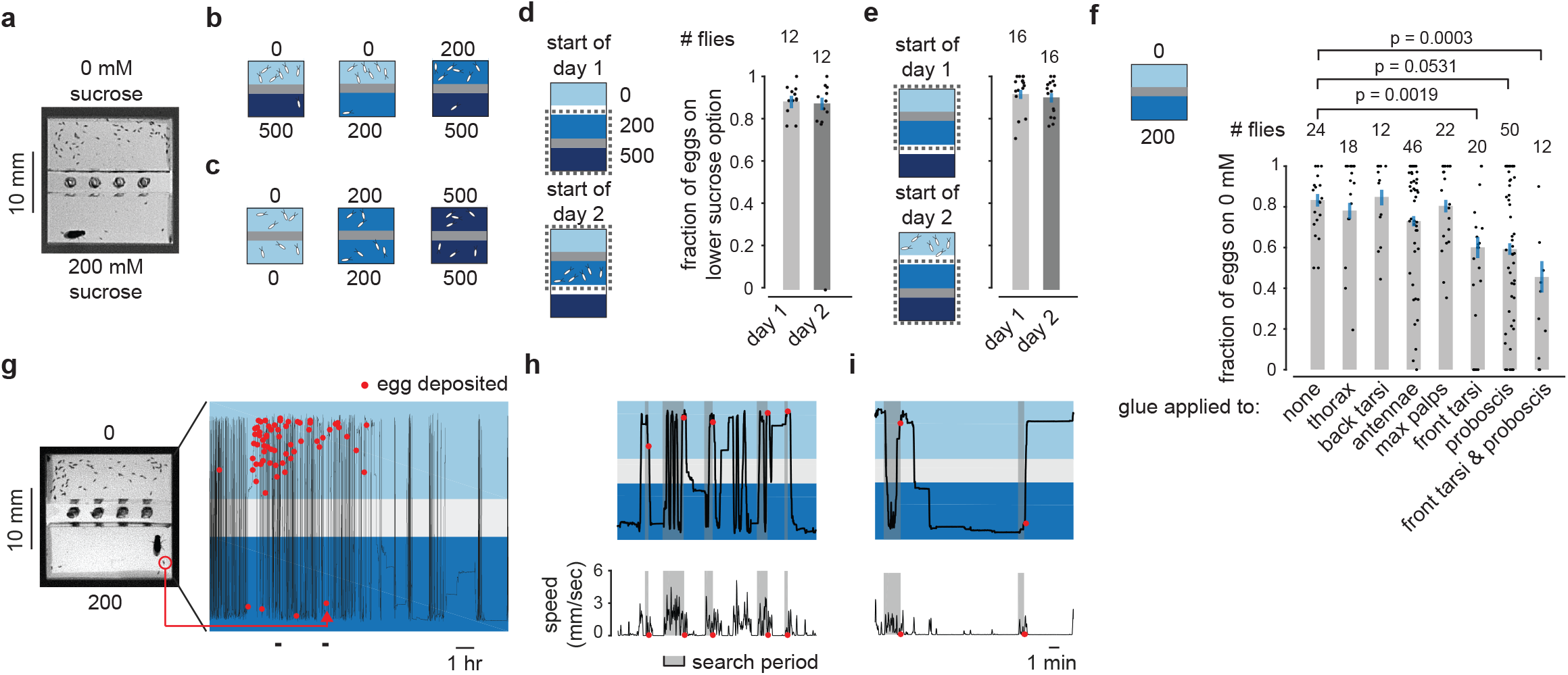
Flies continuously assess the relative sucrose concentration of substrates, using their tarsi and proboscis, to guide egg-laying decisions. **a**, Egg-laying choice chamber with fly. **b, c**, Schematic of relative-value choice data from ref. 13. Each drawn egg represents ∼6 eggs in a real experiment. Numbers indicate sucrose concentration in mM. **d-f**, Fraction of eggs on the lower sucrose option with 95% confidence interval. Each dot represents one fly. In panels d-e, flies laid an average of 40, 42, 42, and 44 eggs per fly, respectively. In panel f, flies laid an average of 25, 24, 29, 25, 29, 18, 23, and 14 eggs per fly, respectively. P-values are 0.0027, 0.0029, and 0.0008 (top to bottom), calculated using a subset of the displayed data (flies that laid ≥ 5 eggs). **g**, Y-position trajectory, and egg-deposition events from a single fly in a high-throughput egg-laying choice chamber. **h**, Zoomed-in view of the first time period indicated at the bottom of the time series in panel g. **i**, Zoomed-in view of the second, later time period indicated at the bottom of the time series in panel g. Locomotor speed in panels h-i is smoothed with a 5 s boxcar filter.

We studied substrates that varied in their sucrose concentrations^5,6^ for two reasons. First, because flies cannot see or smell sucrose at a distance, if flies changed their egg-laying rate on one sucrose substrate due to the existence of a second, we could be sure that this was due to a historical effect of experiencing the second substrate and not due to them currently sensing the distant substrate. Second, previous work has extensively demonstrated that flies make a relative comparison between two sucrose-containing substrates^5,6,13^. That is, if flies are only allowed to lay eggs on 0 mM, 200 mM or 500 mM sucrose substrates, all are acceptable; however, when flies are allowed to choose between any pair of such options, they strongly prefer the lower concentration (schematized in Fig. 1b-c)^13^. Thus, for example, the same 200 mM option is avoided when paired with 0 mM (schematized in Fig. 1b middle) and preferred when paired with 500 mM (schematized in Fig. 1b right).

Prior work has demonstrated that the preference for lower sucrose is not due to flies entirely avoiding the higher sucrose option^6,13^. The preference is also not explained by a competing drive, like feeding, preventing egg laying on higher sucrose, because flies spend a similar amount of time on high and low sucrose prior to laying an egg and we do not observe them to be extending their proboscis (to eat) when walking over the higher sucrose option prior to egg laying^13^. Spatial memory does not seem to serve a prominent role in guiding egg laying in these chambers either^13^. What does seem to explain much of the variance in the flies’ egg-laying choices is their time history of sucrose experiences, as we describe below.

A preference for lower, rather than higher, sucrose may seem counterintuitive because fly larvae (which emerge from the eggs) can metabolize sucrose for energy. However, a soft substrate with relatively low sucrose and high ethanol simulates a rotting fruit^4^, which is the preferred egg-laying environment of most drosophilids^1^. Specifically, the rotting portions of a fruit are relatively depleted of sucrose compared to the ripe parts––due to fermentation of sugars into alcohol––and these regions could be the ideal place for *Drosophila melanogaster* to lay eggs.

### Flies continuously make sucrose assessments of their environment to guide egg-laying decisions

Sucrose-sensing gustatory receptors have been implicated in egg-laying choice^4–6^, suggesting that our flies continuously assessed the local sucrose concentration to guide behavior. However, an alternative possibility is that flies assessed the sucrose concentrations of substrates only at the start of an hours-long experiment and then relied on their initially laid eggs or other chemicals, like deposited pheromones, to guide subsequent egg-laying decisions. Male flies have been shown to deposit pheromones that guide female egg laying^14^, making it plausible that females might do likewise. To test whether substrate marking serves a role in our female-only experiments, we measured egg-laying choice in chambers where we converted a preferred substrate to a non-preferred one, and vice versa, by changing the nature of a second option (Fig. S1b). For example, we let individual flies lay eggs overnight in 200 vs. 500 mM chambers (Fig. 1d left bar). The next day we removed the flies, closed off the 500 mM islands, and opened access to 0 mM islands. We kept the chambers at 4°C to prevent eggs from hatching and then equilibrated the chambers to 24°C prior to assaying another set of flies overnight. Flies laid eggs normally on 0 mM even though 200 mM had many eggs from the start (Fig. 1d right bar). Thus, our flies did not seem to rely on previously laid eggs, or other long-lasting chemical cues, which mark either the preferred (Fig. 1d) or non-preferred (Fig. 1e) option, to guide substrate choice.

Which sensory organs do flies use for assessing the sucrose-content of substrates to guide egg laying? We covered various parts of the fly’s body with light-curable glue^15,16^, likely rendering these parts unable to sense the external chemical environment^17^. We then assessed egg laying in 0 vs. 200 mM chambers. When we applied glue to the front tarsi (front-leg tips) or proboscis, the bias to laying eggs on 0 mM was significantly reduced, and when both these body parts had glue, we could not detect a population-level preference for 0 over 200 mM (Fig. 1f). Choice was not significantly affected, however, when we applied glue to the back tarsi (rear-leg tips), for example, indicating that applying glue, in general, does not affect the ability of flies to target the low sucrose option for egg laying.

Overall, these data argue that flies continuously assess the sucrose content of substrates via receptors on their front tarsi and proboscis to guide egg-laying decisions.

### Two hypotheses

How do flies develop their bias toward laying eggs on the lower sucrose option? One possibility is that flies reduce their probability of laying an egg, or completely turn off egg laying, each time they sense a decrease in sucrose, and vice versa. Another option is that the flies’ substrate experiences over the past few minutes allows them to build an expectation of the sucrose concentrations available in their local environment. This expectation might then allow them to quantitatively adjust their egg-laying probability based on how the sucrose concentration of the current substrate compares to this expectation. In this model, flies would not be relying solely on a comparison with the immediately preceding option. We reasoned that a careful analysis of the spatiotemporal dynamics of egg laying behavior might allow us to differentiate between these hypotheses.

### A framework for understanding how previous substrate experiences guide egg-laying choices

We tracked the x-y position and egg-deposition events of flies in our chambers^13^ and plotted the y-position over time (Fig. 1g-i) (Table S1). A fly was deemed to reside on one of the two substrates based on whether its centroid was above or below the chamber’s midline. Past work that tracked flies during egg laying^6,18^ used chambers in which flies could walk on the hard walls and ceiling (where flies do not lay eggs) and only the x-y position of the centroid was analyzed, making the flies’ precise substrate experiences ambiguous. Because flies could not walk on the ceiling or side walls of our chambers^13^, when their centroid was over a substrate they were physically standing on that substrate.

Prior to laying an egg, flies increase their locomotion during a so-called *search* period^5,13^. The search period begins after ovulation^13^, i.e., after an egg is passed from an ovary to the uterus. It is during search that flies seem to be actively determining on which substrate to lay an egg. To quantitatively define the search epoch preceding each egg, we used a simple algorithm to find the time window prior to each egg-laying event in which the fly’s locomotor activity was elevated^13^ (Fig. 1h-i, gray sections) (Methods).

To examine how a fly’s substrate history impacts egg laying we calculated the fly’s egg-laying rate, during the search period, as a function of time since a substrate transition^13^. We focused on the egg-laying rate during the search period so that periods of time without an ovulated egg would not impact our rate values (e.g., when the fly might be sleeping in our very long experiments). That said, all our conclusions are robust to varying definitions of the search period (Fig. S2). Analyzing hundreds or thousands of eggs was important for accurately estimating rate functions and thus we typically combined data from all tested flies in the same chamber type to generate these curves. We believe that this simplification is reasonable because all combined flies had the same genetic background and flies with outlier behavior were not obvious in visual inspection of the data.

To aid interpretation of the egg-laying rate functions, let us consider some schematic curves from a previous study^13^ (Fig. 2a). A hypothetical fly finishes ovulating and initiates a search on high sucrose 90 s after last experiencing low sucrose. The average egg-laying rate is ∼0.2 eggs/min. (Fig. 2a ex. 1). The fly searches on high sucrose for 30 s and the egg-laying rate is still low, ∼0.2 eggs/min. (Fig. 2a ex. 2). The fly then transitions to low sucrose (Fig. 2a ex. 3). After 15 s, the egg-laying rate is six times higher, ∼1.2 eggs/min. (Fig. 2a ex. 4) and this hypothetical fly deposits an egg. Note that in this analysis, flies make use of substrate experiences regardless of whether these experiences occurred during the current search or not. This interpretation is consistent with the observation that in two-choice sucrose chambers, flies that start a search on higher sucrose know, somehow, to leave that substrate (flies leave higher sucrose in 964 of 1205 or 80% of searches initiated on higher sucrose), whereas flies that start a search on lower sucrose know to stay (flies leave lower sucrose in only 918 of 2744 or 33% of searches initiated on lower sucrose) (p = 3.7e-143)^13^. In addition, note that our egg-laying rate plots do not consider how long a fly has been searching. This simplification is reasonable because flies still lay eggs on low sucrose even after very long searches that have many transitions (Fig. S3). Finally, note that all rate functions, even ones associated with the low-sucrose substrate, start off with a low rate for the first 10 s after a transition. This initially low rate is observed, at least in part, because flies do not lay eggs on the plastic barrier between substrates^13^ and because they are, by definition, walking and not pausing to lay an egg during a transition^13^.

**Fig. 2.**
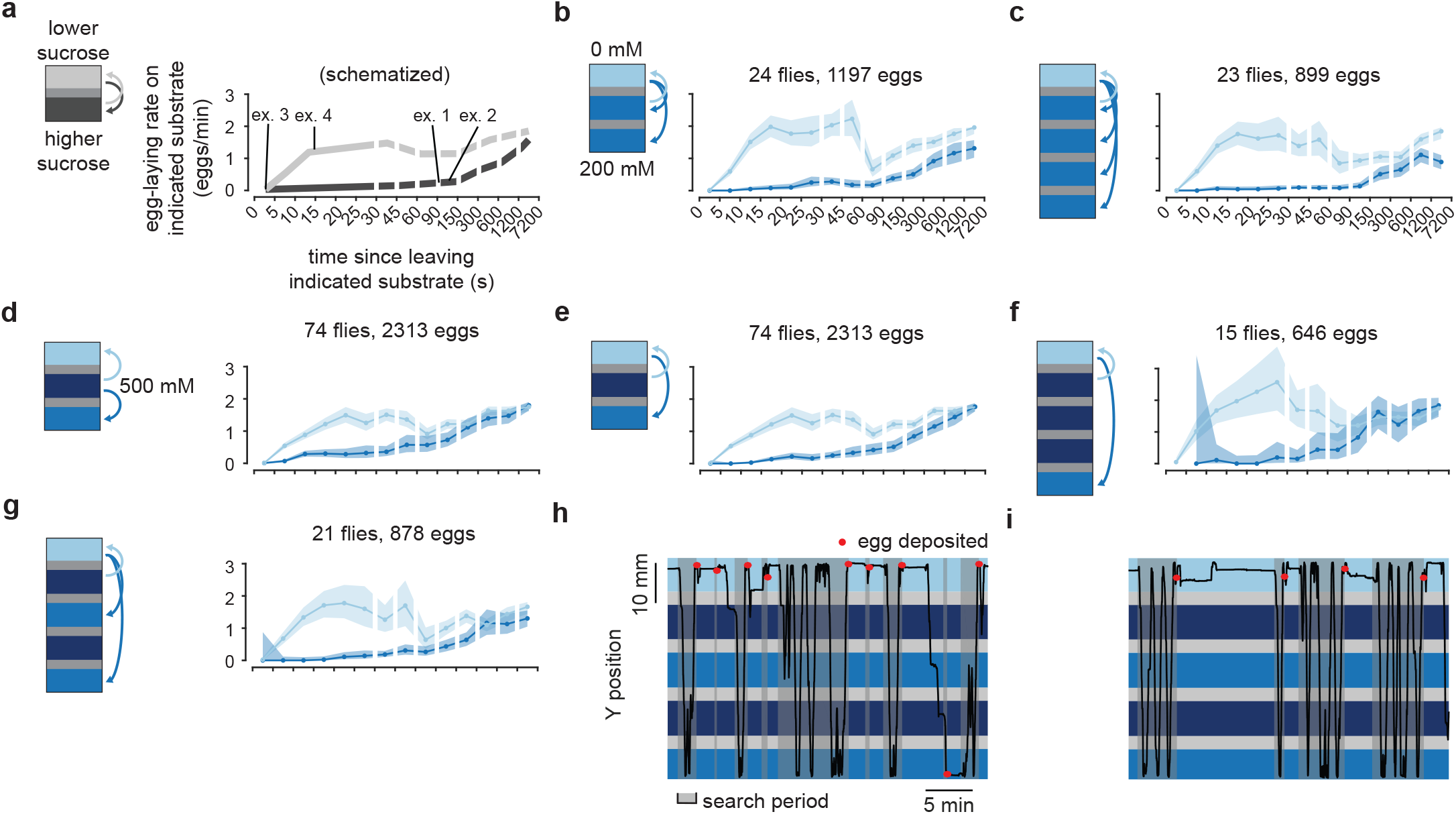
Flies lay most of their eggs on the best of three substrate options. **a**, Schematic of the mean egg-laying rate during the search period as a function of time following a substrate transition (time zero). The rate on each of the two substrates is shown separately (see Main Text and Methods). This schematic is based on data from ref. 13. The light gray curve represents the egg-laying rate on the lower sucrose option as a function of time since last visiting the higher sucrose option (i.e. the transition represented by the light gray arrow on the schematic). Conversely, the dark gray curve represents the egg-laying rate on the higher sucrose option as a function of time since last visiting the lower option. **b-g**, Mean egg-laying rate curves during the search period on a substrate as a function of time since visiting another substrate. 90% confidence intervals shaded. In panels f-g, confidence intervals for 200 mM are large (or missing) for short times because flies rarely (or never) quickly transit between 0 and 200 mM because of the distance between the substrates. **h, i**, Example traces from two separate flies.

Our previously reported result based on these egg-laying rate functions is that when flies transition to high sucrose in two-choice chambers, the egg-laying rate is near 0 for the first ∼2 min. (Fig. 2a darker curve) and then slowly rises to approach the rate on low sucrose^13^ (Fig. 2a darker and lighter curves merge). The fact that the egg laying rates on high and low sucrose become similar over time explains why flies lay eggs similarly in chambers containing only one sucrose option, even when that option is high in sucrose^13^ (Fig. 1c).

### Flies lay eggs on the best of three options

As mentioned earlier, in two-choice chambers flies could, in theory, simply assess whether the sucrose concentration has increased or decreased after each transition. Alternatively, flies might use an internal expectation about substrates. To test between these two hypotheses, we increased the number of substrate options. First, we designed 3 and 5 island chambers (Fig. S1c-d) and confirmed that in these larger chambers flies effectively chose the low sucrose option when presented with two sucrose concentrations, and they did so with similar egg-laying rate dynamics to those observed in standard two-choice chambers (Fig. 2b-c). We then constructed a three-choice chamber with a 500 mM substrate in the middle and 0 mM and 200 mM on either side (Fig. 2d). In this chamber, a fly that lays eggs after any decrease in sucrose concentration should lay eggs upon entering either 0 or 200 mM from 500 mM. However, we observed that flies inhibited egg laying upon entering 200 mM in this chamber (Fig. 2d) (Table S1). As time passed since the last visit to 0 mM, the egg-laying rate on 200 mM gradually increased (Fig. 2e). (The darker blue line in Fig. 2e is the rate on 200 mM since visiting 0 mM, whereas in Fig. 2d it is the rate on 200 mM since visiting 500 mM). These results were robust to increasing the physical distance between the 0 and 200 mM options (Fig. 2f) as well as to chambers in which flies experienced several, locally attractive, increases in acceptability (i.e., decreases in sucrose concentration from 500 to 200 mM) along their path (Fig. 2g-i) (Video S1). In all these three-choice chambers, flies inhibited egg laying nearly completely on 200 mM if they had visited 0 mM in the last ∼30 sec (Fig. 2e-g). This strong inhibition is remarkable because it means that flies were treating 200 mM as a very poor option in these three-choice chambers even though 200 mM is strongly preferred when flies made the identical substrate transition (from 500 to 200 mM) in two-choice chambers^13^ (Fig. 2a). That said, the 0 and 200 mM egg-laying rate functions (Fig. 2e-g) did merge at an earlier time point in three-choice compared to two-choice chambers (Fig. 2a-c), suggesting that some aspect of the three-choice chamber makes differentiating substrates for extended periods more challenging for flies.

### Flies lay eggs more rapidly on high quality substrates that they visit rarely or briefly

If flies can keep track of the option with highest relative value in multi-choice chambers, might they also be able to keep track of how likely they are to experience a good option in a given environment? For example, if a fly rarely experiences a 0 mM substrate, would it lay an egg more quickly when it does experience it, consistent with the fly being positively surprised?

We designed chambers where flies would experience 200 mM for a long time before entering 0 mM (Fig. S1e-g). In chambers with a tiny (2 mm) tunnel between substrates, flies laid eggs at a higher rate in the 5-10 s window after a transition to 0 mM compared to very similar chambers with a 4 mm tunnel (Fig. 3a-b bin indicated with arrow). To further restrict flies from entering the 2 mm tunnel, we designed a chamber with angled walls near the tunnel to “bounce” flies that are walking along the edge of the chamber away from the tunnel. In this chamber, flies laid eggs even more rapidly upon entering the 0 mM side (Fig. 3c bin indicated with arrow) (Video S2).

**Fig. 3.**
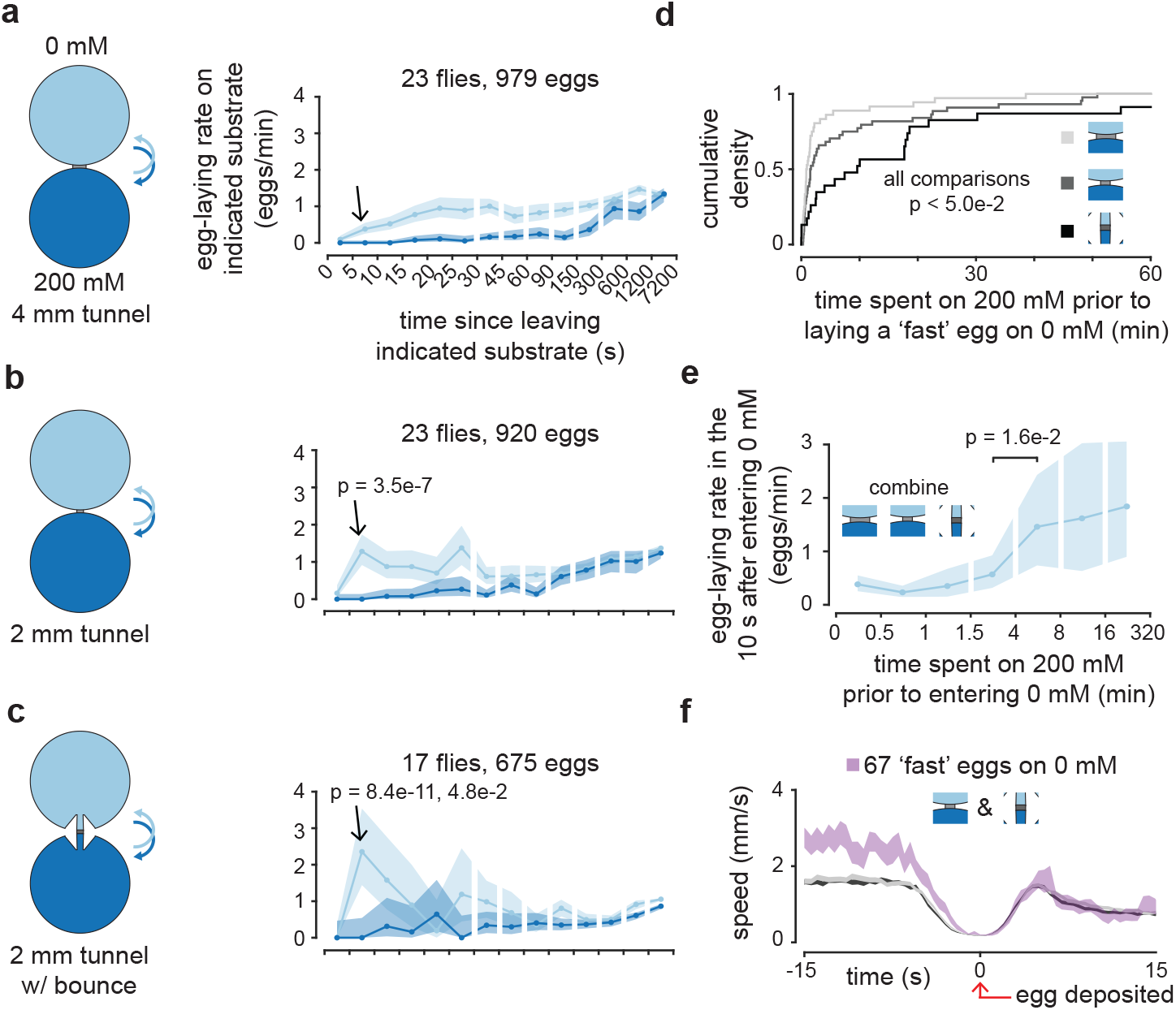
Flies lay eggs faster when on a substrate that they have not experienced recently. **a-c**, Mean egg-laying rate during the search period on a substrate as a function of time since visiting another substrate. 90% confidence interval shaded. P-value in panel b is comparing the indicated bin to the same bin in panel a. P-value in panel c is comparing the indicated bin to the same bin in panels a and b, respectively. **d**, Cumulative distribution of time spent on 200 mM prior to entering 0 mM and laying an egg in 10 s or less (i.e., ‘fast’ eggs). Medians are 54, 96, 576 s for increasing shades of gray. 36, 44, 23 eggs for increasing shades of gray. **e**, Mean egg-laying rate during the search period in the 10 s after entering 0 mM as a function of time previously spent on 200 mM. 90% confidence interval shaded. **f**, Mean locomotor speed aligned to egg deposition. Data in gray is all eggs from the chambers as colored in panel d. For increasing shades of gray, we analyzed the following number of eggs: 979, 918 (two less than panel b because eggs that were laid in the first or last few minutes of a video were not included in all triggered averages to prevent averaging of partial traces), and 675.

Do flies lay eggs quickly on the 0 mM substrate because they have experienced the relatively-worse 200 mM substrate for a long time? As expected, in the chambers shown in Figure 3a, Figure 3b and Figure 3c respectively, flies tended to spend progressively more time on the 200 mM substrate before laying ‘fast’ eggs (eggs laid within 10 s of entering 0 mM) (Fig. 3d). Similarly, when we combine all the egg-laying events from these three chambers, we observe that flies increase their egg-laying rate upon entering 0 mM specifically if they have not experienced 0 mM for > 4 minutes (Fig. 3e). The time delay between the dip in the flies’ locomotor speed and egg deposition is similar for fast eggs (Fig. 3f pink) and all eggs in these three chambers (Fig. 3f shades of gray), suggesting that the decision to lay an egg, and not the egg-deposition motor program is made faster. Note that the absolute locomotor speed prior to egg laying is higher for fast eggs because flies need to be moving to cross the barrier between substrates. These results suggest that flies make faster egg-laying decisions when they find an unexpected, preferred substrate.

We noticed that flies often circled our chambers while staying adjacent to the walls (thigmotaxis), which meant that they had shorter transit times through the middle islands of rectangular chambers (median, ∼4.5 s) compared to the edge islands (median, ∼13 s) (Fig. 4a). We thus posited that if the preferred substrate were in the briefly visited middle island, flies may need to make a faster decision to lay an egg on that substrate. We found that flies indeed lay eggs more quickly after a transition to the preferred substrate in these chambers (Fig. 4b bin indicated with arrow) compared to chambers where the preferred substrate was on either edge (Fig. 2a-g). This result was clear in individual trajectories (Fig. 4c) (Video S3). Interestingly, concomitant with faster egg laying on the middle substrate, flies also had a higher egg-laying rate than expected (from our other three-choice experiments) on the closest alternative (compare medium blue curve from 0 to 90 s in Fig. 4d with Fig. 2e). Such anomalous eggs on an intermediate option may be undesirable and minimizing them may be one of the reasons why flies resort to quick egg laying only when necessary (i.e., when a good substrate is experienced only briefly) (see Discussion).

**Fig. 4.**
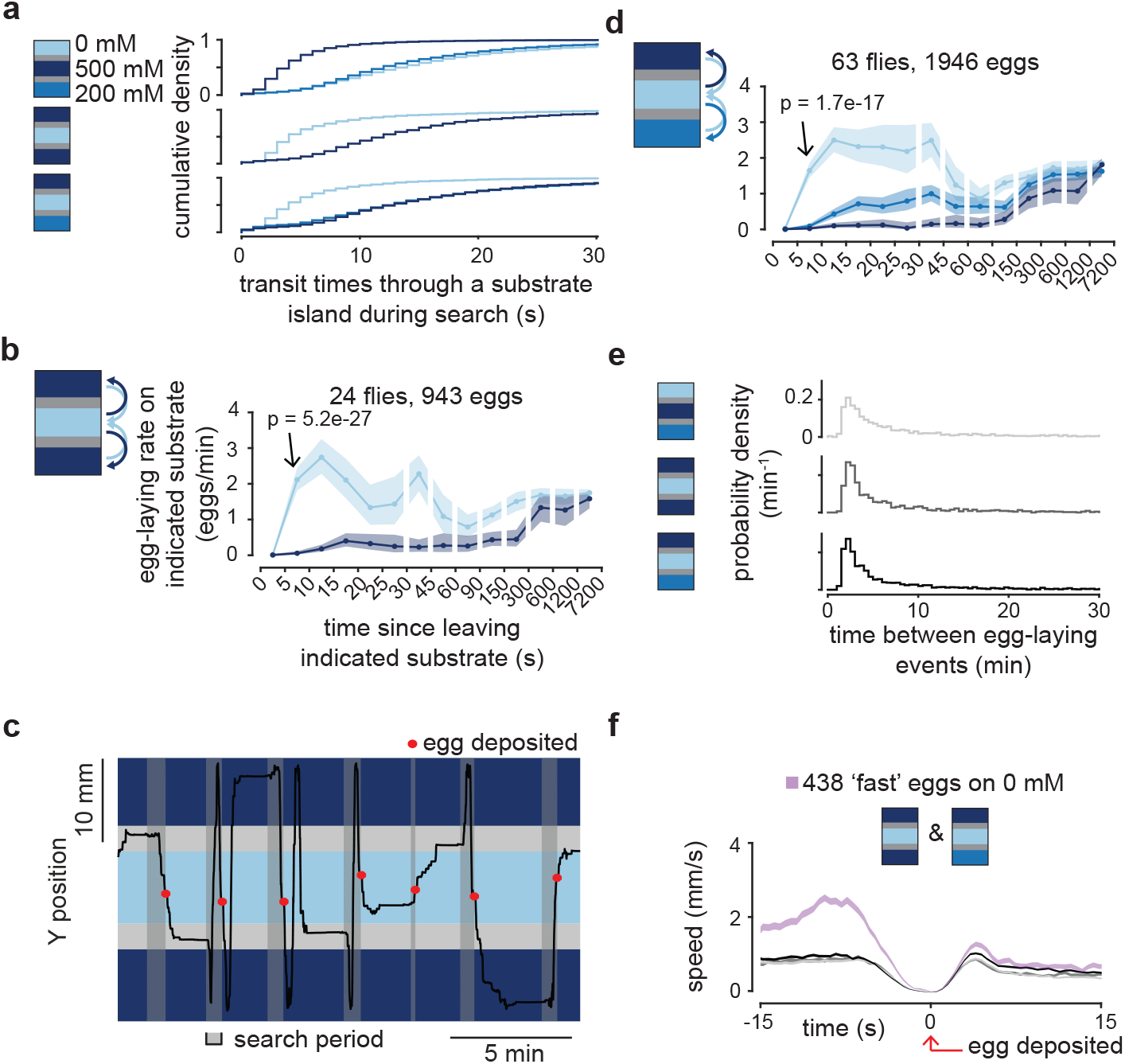
Flies lay eggs faster when on a substrate that they typically visit briefly. **a**, Cumulative distribution of transit times through substrate islands during the search period. 2637, 5694, and 2903 transit times for 0, 500, and 200 mM, respectively, in 0/500/200 chamber. 1625 and 1508 transit times for 0 and 500 mM, respectively, in 500/0/500 chamber. 1741, 3354, and 1810 transit times for 500, 0 and 200 mM, respectively, in 500/0/200 chamber. **b**, Mean egg-laying rate during the search period on a substrate as a function of time since visiting another substrate. 90% confidence interval shaded. P-value is comparing the indicated bin to the same bin in Figure 2b. P-value is 1.4e-12 if compared to a 0/500/500 mM chamber (Table S1). **c**, Example trace. **d**, Mean egg-laying rate as in panel b. **e**, Time between egg-laying events. Medians are 5.1, 4.1, and 4.4 min, respectively. 2262, 922, and 1901 inter-egg intervals, respectively. **f**, Mean locomotor speed aligned to egg deposition. Data in gray is all eggs from the chambers as colored in panel e. 2312, 943, 1946 eggs for increasing shades of gray.

Importantly, the inter-egg interval is not increased when the preferred substrate is in the middle island suggesting that flies are not abandoning egg-laying search attempts in these chambers (which would artificially increase egg-laying rates if these abandoned search attempts are not included in our search period definition) (Fig. 4e). Furthermore, the increased egg-laying rate with briefly visited substrates holds for broader definitions of the search period (Fig. S2e). As in our previous experiments, the egg-deposition motor program (as proxied by the dip in locomotor speed) is not accelerated (Fig. 4f). Flies thus decide to lay eggs faster on the best option in environments where they typically visit that option only briefly.

Together, these data support the hypothesis that flies build an internal expectation regarding their substrate environment and use this expectation to guide egg-laying decisions. This expectation keeps track of the best option experienced over the past few minutes (Fig. 2) alongside of how rarely or briefly a substrate is experienced (Figs. 3-4).

### Different *Drosophila* strains and species show different egg-laying rates in the same environment

How flies interpret substrate experiences for egg laying may differ depending on the needs and preferences of specific strains or species. We tested two other wild type strains of *Drosophila melanogaster* in 0 vs. 200 mM chambers and found that both released inhibition of egg laying on 200 mM faster than did Canton-S (CS) flies (Fig. 5a-b compared to Fig. S2a). For example, the TUT strain showed no statistically detectable evidence of substrate history after only ∼15 s of being on a new substrate (Fig. 5b). One interpretation is that each wild type strain readjusts its expectations on a timescale that is tuned to the statistics of the niche from which it was isolated. For example, flies that tend to experience frequent fluctuations in substrate quality in their natural environment may readjust their expectations more slowly– –i.e., hold out longer for a better option––than those that do not. Alternatively, certain fly strains may be more capable than others at building or using expectations due to genetic constraints whose nature is not yet clear.

**Fig. 5.**
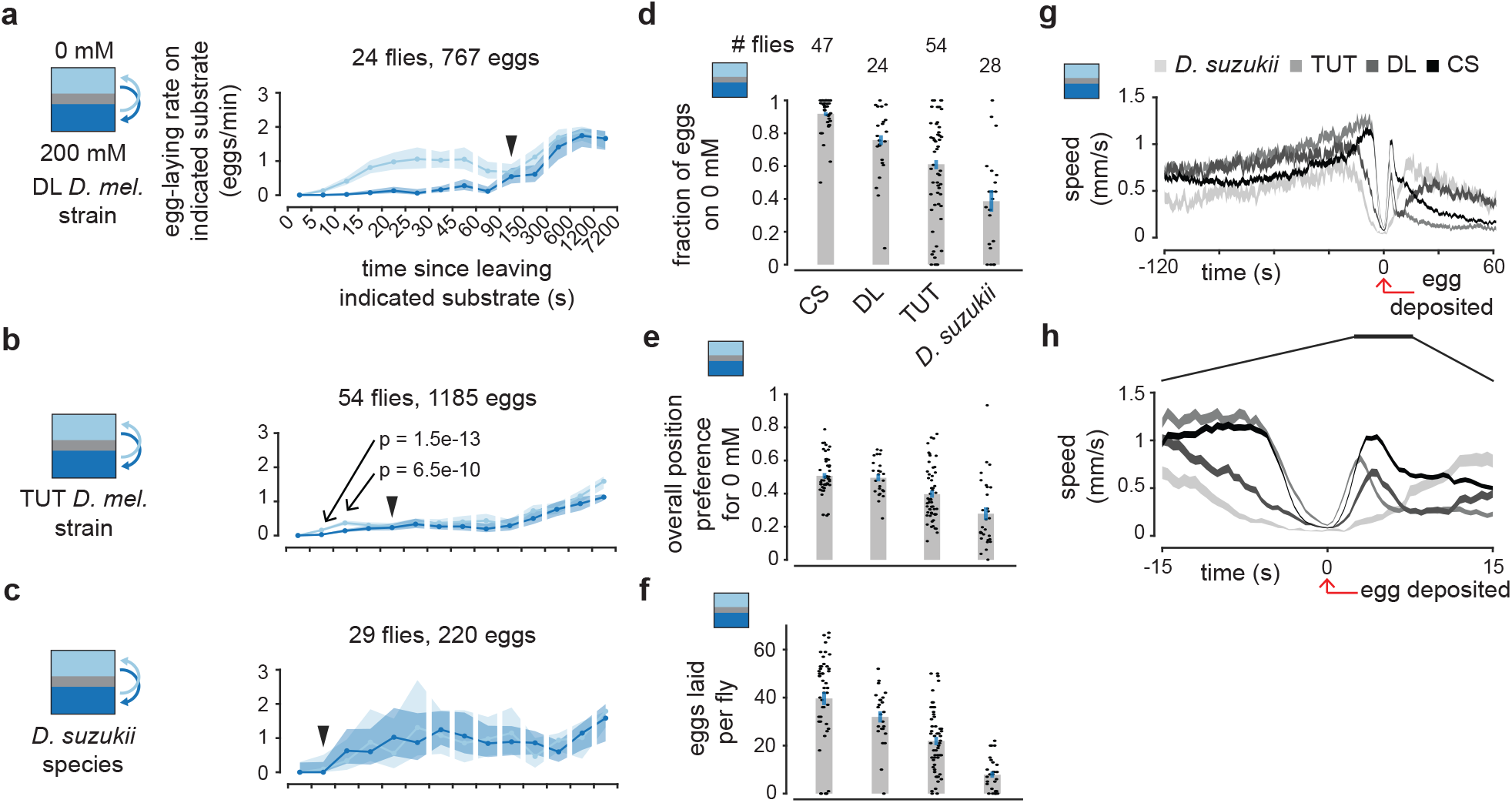
Different *Drosophila* strains and species show different egg-laying rates in the same environment. **a-c**, Mean egg-laying rate during the search period on a substrate as a function of time since visiting another substrate. 90% confidence interval shaded. P-values are comparing 0 and 200 mM for the indicated bin. Filled black arrows indicate first bin past 5 s for which p > 0.05 (i.e., a quantitative estimate of when the curves merge). **d**, Fraction of eggs on 0 mM with 95% confidence interval. Each dot represents one fly. **e**, Fraction of time spent on 0 mM per fly. Each dot represents one fly. **f**, Eggs laid per fly. Each dot represents one fly. **g, h**, Mean locomotor speed aligned to egg deposition. 214, 1184, 766, and 1862 eggs for increasing shades of gray.

Most species of *Drosophila* prefer to lay eggs on rotten fruit^1^. However, *Drosophila suzukii* prefer to lay eggs on ripe fruit^1,19^. A variety of evolutionary changes have enabled this switch in *D. suzukii*, including an enlarged and serrated ovipositor for penetrating ripe fruit^19^. We found that unlike the three wild type strains of *D. melanogaster, D. suzukii* did not exhibit a measurable egg-laying preference for lower sucrose substrates in our paradigm, i.e., the egg-laying rates on 0 and 200 mM were similar in a two-choice chamber (Fig. 5c). *D. suzukii* actually laid slightly more eggs on the higher (200 mM) sucrose substrate (Fig. 5d) consistent with their overall preference toward spending more time on high sucrose (Fig. 5e). A loss of preference for low sucrose may be an adaptation that helps *D. suzukii* lay eggs on ripe fruit. In general, different strains/species laid a different number of eggs (Fig. 5f) which may be indicative of their egg laying tendencies in the wild or their sensitivity to our experimental conditions. All strains/species, on average, increase their locomotor speed (search) prior to pausing to deposit an egg (Fig. 5g). However, each strain/species has a different time delay between the dip in average locomotor speed and egg deposition (Fig. 5h). It is possible, for example, that *D. suzukii* has a longer egg-deposition motor program aimed to penetrate harder, ripe fruit.

These results suggest that aspects of the expectation processes may be evolving with the needs and preferences of different *Drosophila*.

### Dopa decarboxylase expressing neurons are a strong candidate for performing expectation-related calculations

We screened for neurons whose electrical activity contributes to normal 0 vs. 200 mM egg-laying choice. We reduced the excitability of genetically targeted neurons by expressing in them, via the Gal4/UAS system, the Kir2.1 potassium channel^20^. A temperature-sensitive Gal80 transgene^21^ was used to restrict expression of *kir2*.*1* until a day before the assay. Control and experimental flies were siblings with the identical genotype, but experimental flies were held at 31°C, instead of 18°C for controls, for a 23-hour period prior to the egg-laying assay, during which *kir2*.*1* gene expression was permitted. Egg-laying assays for all genotypes, experimental and controls, were performed at the same, 24°C, temperature (Methods). The Gal4 lines tested in this screen were chosen either (1) randomly, (2) based on previous egg-laying studies^5–7,22^, or (3) based on hits identified earlier in the screen. Of the 115 Gal4 lines tested, in the control condition 110 (96%) laid more eggs on 0 than 200 mM (Fig. S4a) and 111 (97%) laid 5 or more eggs per fly (Fig. S4b). To visualize the results of the screen, we plotted the p-value versus fold-change-in-choice (experimental choice divided by control choice) (Fig. 6a). Remarkably, of the 115 Gal4 lines tested, the top 8 hits, and many of the hits just outside of the top 8, had the Gal4 transgene driven by a Dopa decarboxylase (Ddc) related enhancer (Table S2). We performed more detailed experiments on the DDC-Gal4 line^23^, specifically, to see if we could better understand how these neurons contribute to egg-laying choice.

**Fig. 6.**
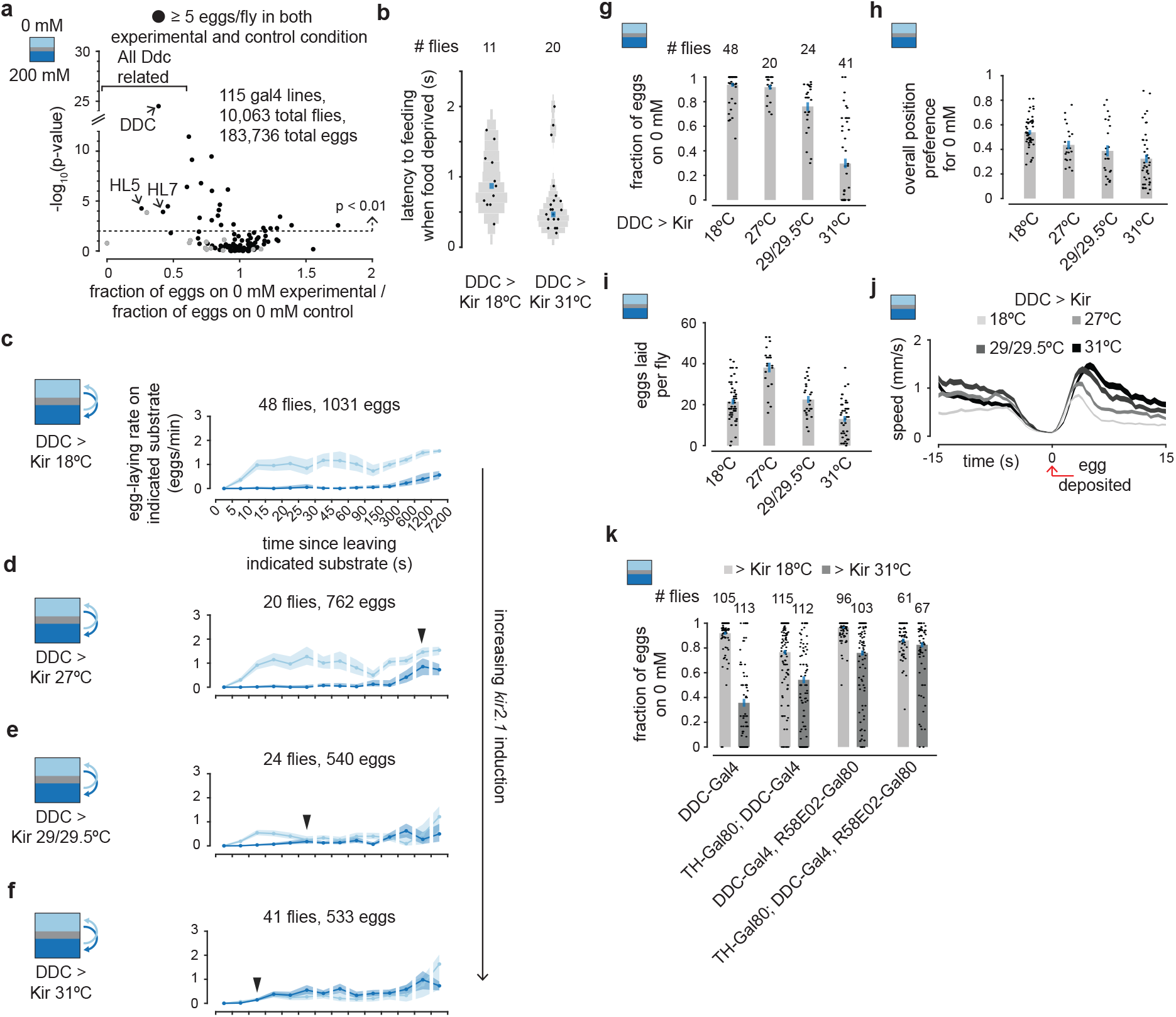
Dopamine neurons are a strong candidate for performing expectation-related calculations. **a**, Scatter plot of individual Gal4 lines assayed in our egg-laying decision screen. The x-axis represents the fraction of eggs laid on 0 mM, after pooling all eggs from a given experimental genotype, divided by the same value for control flies. The y-axis shows the p-value for the difference between the experimental and control groups. The top 3 Gal4 lines with ≥ 5 eggs per fly in both the experimental and control condition are labeled. **b**, Latency to feeding by food deprived flies after crossing plastic barrier from 0 mM to 200 mM. **c-f**, Mean egg-laying rate during the search period on a substrate as a function of time since visiting another substrate. 90% confidence interval shaded. Filled black arrows indicate first bin past 5 s for which p > 0.05 (i.e., a quantitative estimate of when the curves merge). **g**, Fraction of eggs on 0 mM with 95% confidence interval. Each dot represents one fly. **h**, Fraction of time spent on 0 mM per fly. Each dot represents one fly. **i**, Eggs laid per fly. Each dot represents one fly. **j**, Mean locomotor speed aligned to egg deposition. 1029, 761, 540, and 533 eggs for increasing shades of gray. **k**, Fraction of eggs on 0 mM with 95% confidence interval. Each dot represents one fly. In panels a-k, all flies have tubulin promoter driving Gal80^ts^.

First, as a control, we checked if DDC-Gal4 experimental flies––hereafter, DDC > Kir (31°C pre)––could still sense sucrose. We assayed proboscis extension of food-deprived flies in 0 vs. 200 mM chambers with a movable barrier between substrates (Fig. S1h, Methods). Flies that were not food deprived, regardless of pretreatment, showed no measurable proboscis extension upon entering 200 mM (data not shown). DDC > Kir (31°C pre) food-deprived flies either extended their proboscis or, in cases where our camera could not see their proboscis clearly, entered an obvious feeding posture quickly after crossing onto 200 mM (Fig. 6b) (Fig. S5). Thus, DDC > Kir (31°C pre) flies can still sense the sucrose content of a substrate, suggesting that the role of Ddc neurons is downstream of sensory processing.

We induced *kir2*.*1* at four different levels in the DDC-Gal4 neurons, using four temperatures of pretreatment to remove varying amounts of the Gal80^ts^ inhibition (Fig. 6c-f). The more Kir2.1 we expressed, the earlier in time after a substrate transition did the 0 and 200 mM egg-laying rate curves begin to merge (Fig. 6c-f, filled black arrows). When Kir2.1 expression was strongest in DDC > Kir (31°C pre) flies, the egg-laying rate curves were nearly superimposable throughout (Fig. 6f) meaning that these flies laid eggs similarly on 0 and 200 mM substrates. Importantly, the DDC > Kir phenotype is different from one where flies shift preference to sucrose or generally increase variability in behavior because, in these two interpretations, the 200 mM egg-laying rate curve should increase equally in all bins regardless of the time since a transition. Instead, flies show properly biased egg-laying rates proximal to a substrate transition in the two intermediate pretreatment temperatures (27°C and 29/29.5°C) (Fig. 6d-e). We interpret these results to mean that expectation-related information––either the value of the expectation and/or the comparison of this signal with the value of the current substrate (i.e., a relative-value signal)––is degraded when Ddc neurons are made leakier with Kir2.1. The overall egg-laying rate during search tended to drop as Kir2.1 expression was increased (progressively lower mean curve levels in Fig. 6d-f), meaning that flies searched for longer prior to laying an egg. This overall drop in the egg-laying rate is thus also consistent with the notion that relative-value signals are compromised with Kir2.1 expression. As expected from the egg-laying rate curves, we found that increased induction of *kir2*.*1* resulted in a progressive decrease in the fraction of eggs laid on 0 mM (Fig. 6g). The fact that DDC > Kir (31°C pre) flies laid more eggs on 200 mM than 0 mM is consistent with them spending more time, overall, on the 200 mM side (Fig. 6h) alongside their indifference as to where to lay eggs (Fig. 6f). The decrease in egg-laying preference of DDC > Kir (31°C pre) flies (Fig. 6c-f) was unrelated to the number of eggs laid (Fig. 6i) or to changes in the egg-deposition motor program (as proxied by the dip in locomotor speed) (Fig. 6j).

### Dopamine neurons may be the relevant set within the Dopa decarboxylase expressing population

The Dopa decarboxylase enzyme is involved in the biosynthesis of both dopamine and serotonin. Neurons in which the enhancer for Dopa decarboxylase is active are found in the *Drosophila* brain and ventral nerve cord (Fig. S6). In vertebrates, certain populations of dopaminergic neurons are involved in expectation-related calculations^24–29^. To test if the dopamine neurons, specifically, within the DDC-Gal4 population were important for choice, we used TH-Gal80^30^ (tyrosine hydroxylase related enhancer) and/or R58E02-Gal80^31^ (dopamine transporter related enhancer) transgenes to minimize or eliminate Kir2.1 expression in the dopaminergic subset of cells targeted in the DDC-Gal4 line. Expression of Gal80 by either of the abovementioned enhancers partially rescued the choice defect in DDC > Kir (31°C pre) flies. With both enhancers driving Gal80 the choice defect was fully rescued (Fig. 6k). These results support the hypothesis that it is the dopamine neurons within the set of cells targeted in the DDC-Gal4 line whose normal electrical activity is relevant for choice. The fact that TH-Gal80 or R58E02-Gal80 alone does not rescue the choice defect completely suggests that many dopamine neurons may be involved. However, it is formally possible that each Gal80 transgene progressively decreases Kir expression in the critical, small subset of DDC-Gal4 neurons.

## Discussion

We found that *Drosophila* egg-laying decisions are impacted by past substrate experiences in a non-trivial manner. We argue that the substrate-history effects described here can be understood via a framework in which flies internally construct an expectation of the substrate composition of their environment. This internal expectation is then compared to the quality of the current substrate to guide egg-laying decisions.

### A model for expectation-guided egg laying

A set of descending neurons, called *oviDNs* (oviposition descending neurons), are required for laying eggs^32^. We have recently imaged calcium signals in oviDNs and their activity dips during ovulation, rises over many seconds to minutes during search, and crosses a threshold level immediately prior to the final abdomen bend for egg laying^13^. The rate of rise of the oviDN calcium signal during search is modulated by the relative value of the current substrate. We hypothesize that an internal expectation signal––inferred to exist from the experiments discussed here––is compared to an estimate of the value of the current substrate, with the difference between these two variables governing the oviDN signal’s rate of rise and thus its likelihood to hit threshold (Fig. 7). For example, when a fly is surprised by an unexpected, preferred substrate, the difference in value between the current substrate and the expectation is large and, as a result, the slope of the oviDN calcium rise is large. A large slope causes the oviDNs to hit threshold quickly, resulting in a fast decision to lay an egg. Future neurophysiological experiments should explore the activity of oviDNs in such conditions.

**Fig. 7.**
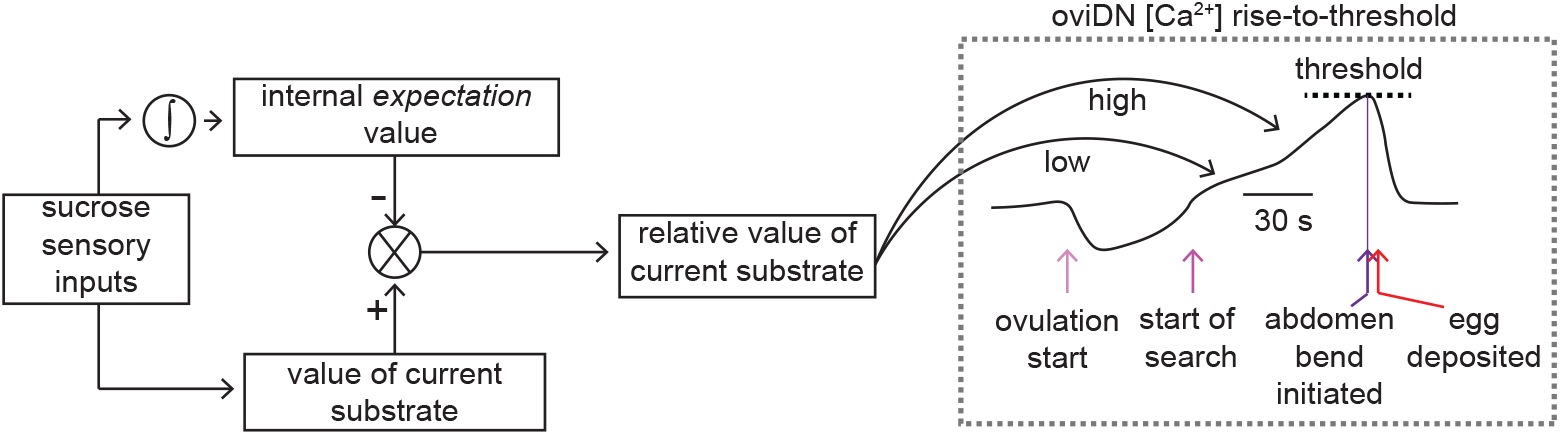
A model for expectation-guided egg laying. The integral symbol indicates the brain integrating sucrose sensory inputs over time to build an expectation signal. The *expectation* value signal is subtracted from the value of the current substrate to create a relative value signal. This relative value signal regulates a rise to threshold process to drive egg laying.

### An internal sense of the external environment that guides behavior

An internally constructed expectation that allows an animal to change its response to the same stimulus based on an informed prediction is a signature of cognitive behavior^33^. Indeed, one way to conceptualize our findings is that flies latently learn^34,35^ (i.e., learn without obvious positive or negative reinforcement) an internal representation of their external environment, which shapes responses to future stimuli. Several studies in *Drosophila* have shown that adult flies and larvae use sensory experiences over the past few seconds, which were not obviously reinforced, to regulate behavior, including during courtship^36^, taxis^37^, and spatial orientation^38^. Similar short-timescale comparisons have been observed even in *E. coli* during chemotaxis^39,40^. The longer, minute-timescale behavioral changes we observe here suggest that historical information may be converted into an explicit, stable internal expectation signal in the fly brain, which future neurophysiological work should aim to reveal.

What form might an internal expectation take? One possibility is that flies build an explicit mental model of their environment based on a remembered list of the substrates they previously visited, augmented with the times these substrates were encountered and the durations spent on each. A perhaps more realistic possibility––given the small size of the *Drosophila* brain––is that flies convert their substrate experiences into an analog neural signal, or a set of such signals, which encode an expectation value of the environment. For example, a single neuron could track this *expectation* explicitly via its spike rate, built off recent substrate experiences. In a 0 vs. 500 vs. 200 mM chamber (Fig. 2e), this neuron’s spike rate would be continuously high (assuming the fly has recently visited 0 mM). The egg-laying rate would thus be low unless the fly is situated on 0 mM, which is the only substrate that would be able to drive neurons tracking the value of the current substrate to spike at a high enough rate to exceed the high rate of the expectation neuron (Fig. 7). If the same fly were transferred to a 500 vs. 0 vs. 200 mM chamber (Fig. 4d), the expectation-neuron’s spike rate would drop because the best option is experienced more briefly. This lower spike rate would make the *current*-minus-*expectation* output high enough on both 0 and 200 mM to cause an increase in the egg-laying probability on both options, explaining the faster egg laying on 0 mM and the increased, anomalous eggs on the closest alternative. Note that this putative expectation-neuron’s spike rate may need to track more than just the mean sucrose concentration experienced over the past few minutes because if flies were merely using such a mean signal they would, for example, have had a hard time inhibiting egg laying on the 200 mM option in 0 vs. 500 vs. 500 vs. 500 vs. 200 mM chambers (Fig. 2f). Although a single expectation neuron is possible, it is important to note that our experiments point to a wider array of neurons being involved in the expectation process (Fig. 6k). Whereas our work here focused on sucrose-mediated choice, future work should explore whether this internal expectation incorporates sensory modalities other than sucrose and if different modalities have independent expectation signals.

### Dopamine neurons and expectations

Monoaminergic neurons have been implicated in increasing or decreasing the fraction of eggs laid on a preferred substrate^6,7,22^. However, this past work did not provide a computational framework with which to interpret the role of these neurons in egg-laying decisions. A clear hypothesis for the computations involved in egg-laying substrate choice (Fig. 7) will be invaluable for interpreting neurophysiological signals^41^, especially if the relevant signals turn out to be distributed across a broad set of neurons, as our data suggest (Fig. 6k).

A recent report has argued that Parkinson’s disease patients have impairments in their ability to use priors (i.e., expectations) for decision making^42^, which is reminiscent of the behavioral phenotype we observe when Ddc expressing neurons are inhibited in egg-laying *Drosophila*. Parkinsonian patients suffer from a progressive loss of midbrain dopamine neurons. In vertebrates, at least some dopamine neurons signal a reward prediction error^24–26^, which represents the difference between the value of a current experience and the expectation of value associated with that experience. It is believed that these reward prediction error signals act both to update expectations about the world and also to modify ongoing behavior^27–29^. A reward prediction error is akin to the proposed relative-value output signal in our model (Fig. 7). If dopamine neurons carried such a signal in egg-laying flies––and thus modified either the fly’s expectations about the world (pathway not diagrammed in Fig. 7) or the rate-of-rise of the oviDN signal to more directly impact ongoing action––it would make sense why their activity is critical for egg-laying substrate choice. Dopamine signals in *Drosophila* have been shown to track the value of certain states in a seemingly absolute manner^31,43,44^ and it is possible that future neurophysiological work will reveal that fly dopamine cells also signal value in a relative manner^45^.

Although the physiology of dopamine neurons has been well studied in vertebrates^24–29^, many questions remain^46^. For example, little is known about how expectations are calculated to create reward-prediction error signals^47–49^ or how the output of dopamine neurons modulates action^50,51^. Our work suggests that a deeper understanding of the neural basis of egg-laying decisions in the tractable^52–54^ fly brain may help to reveal general principles related to value signaling and dopamine physiology which have proven harder to identify in larger nervous systems.

## Supplement

### Supplemental Figure Legends

**Fig. S1.**
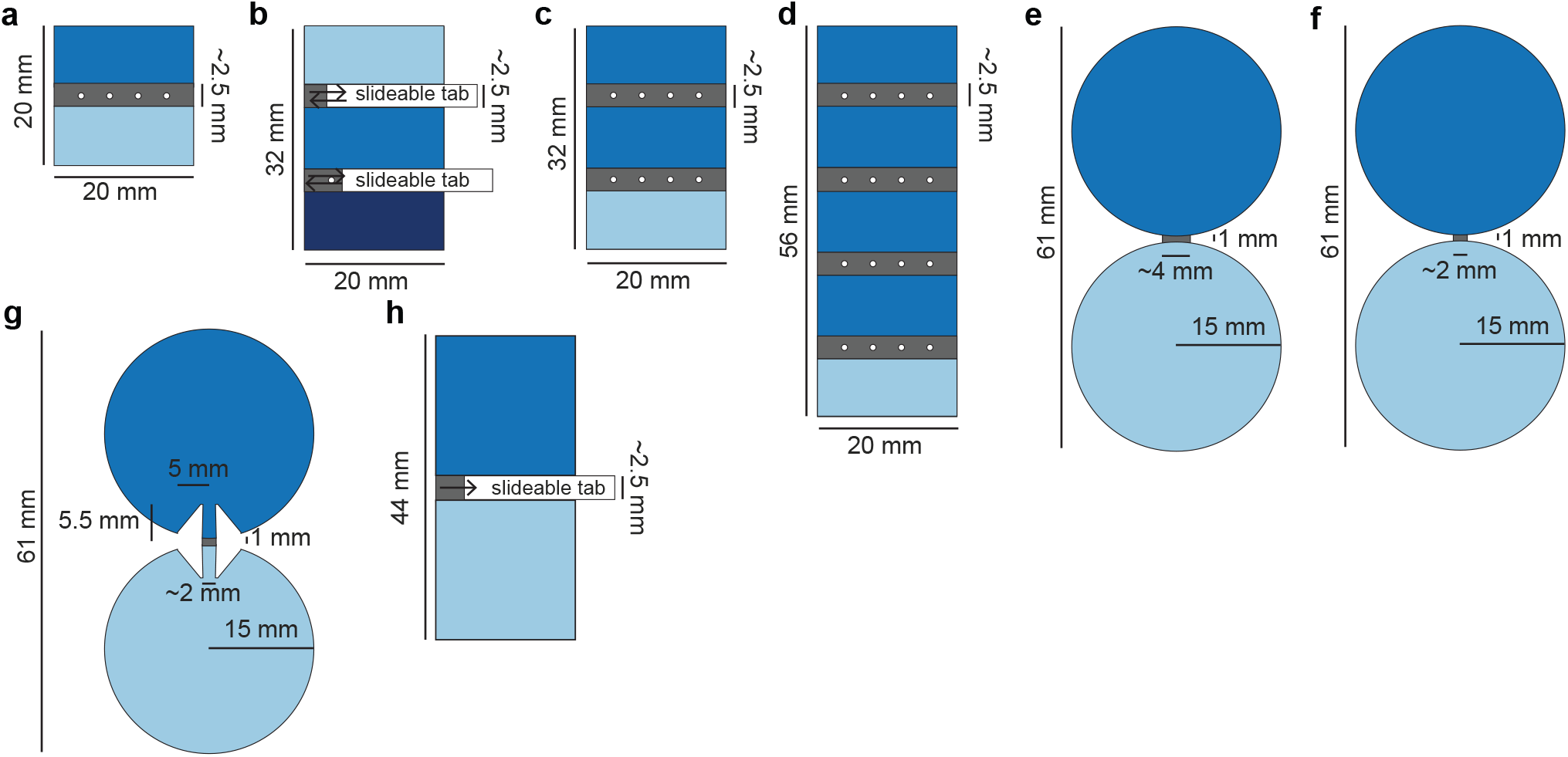
Egg-laying choice chambers used in this study, related to Figures 1 through 6. **a**, Two-island chamber. **b**, Three-island chamber with movable boundaries used in Fig. 1d-e. **c**, Three-island chamber. **d**, Five-island chamber. **e-g**, restricted corridor (and control) chambers used in Fig. 3a-f. **h**, Movable boundary chamber used in Fig. 6b.

**Fig. S2.**
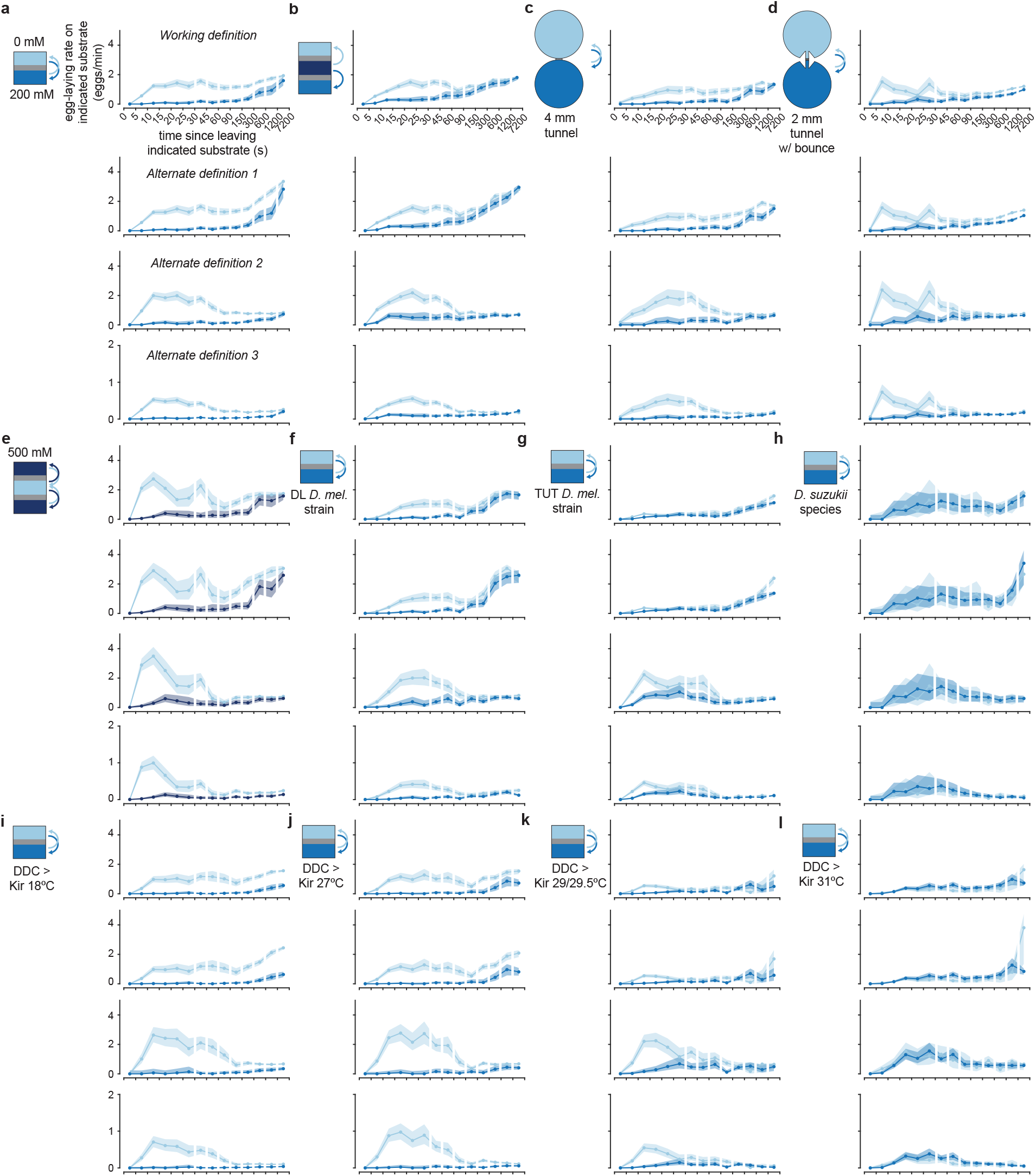
Egg-laying rate functions with different definitions of the *search* period, related to Figures 2 through 6. **a-l**, Egg-laying rate functions in twelve chambers with different search-period definitions. *Working definition*: Egg-laying rate functions as defined in this study and a previous study^13^ (Methods). *Alternate definition 1*: Egg-laying rate functions with search durations under 30 s not set to 30 s. *Alternate definition 2*: Egg-laying rate functions with the search period always set to the 90 s before each egg-laying event. If this 90 s extends into the previous egg-deposition event plus 1 minute, it was shortened. This prevents search periods from overlapping with other search periods or with ovulation (which takes at least 1 minute)^13^. *Alternate definition 3*: Same as *alternate definition 2* except we changed 90 s to 15 min.

**Fig. S3.**
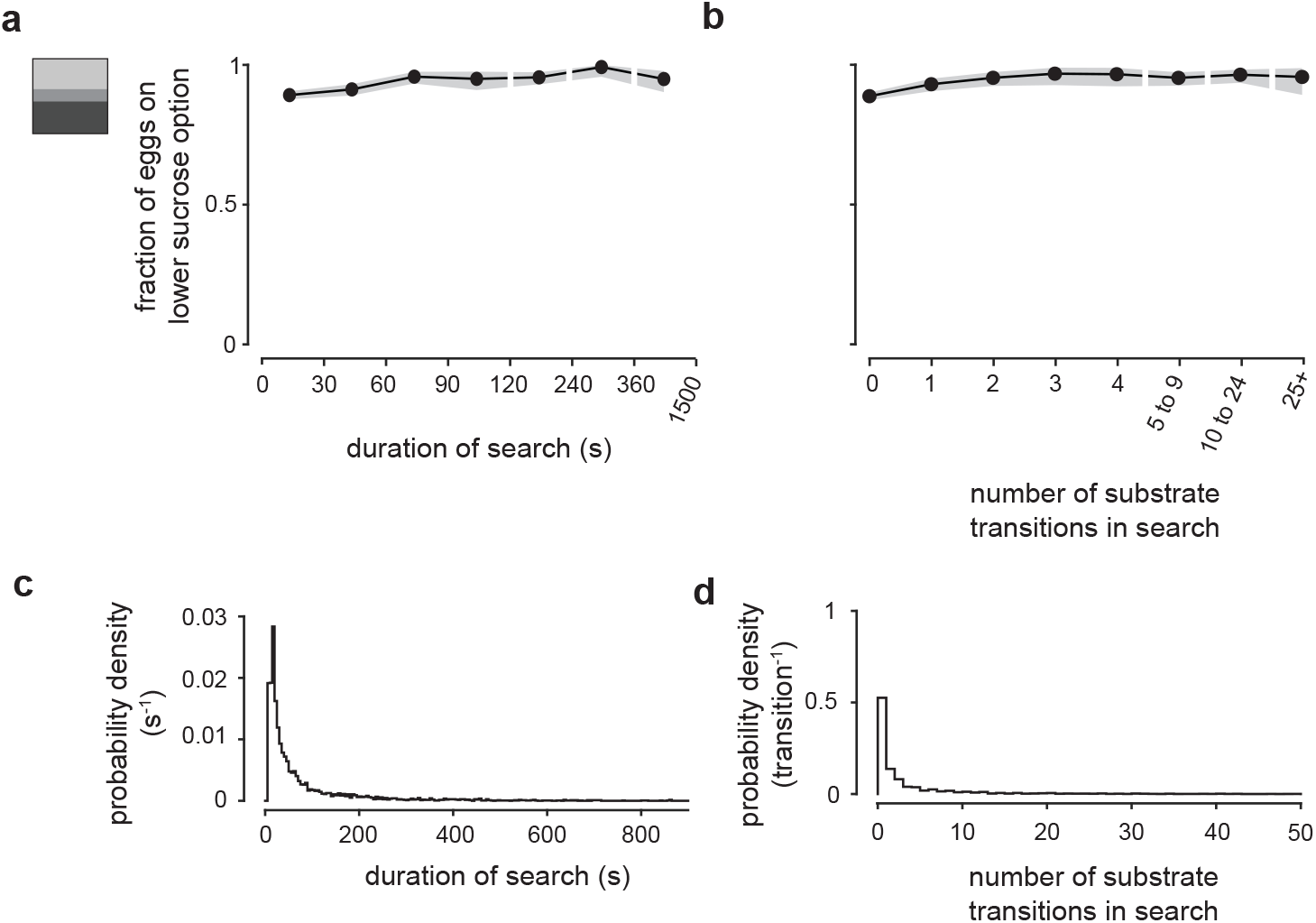
Egg-laying behavior as a function of search duration, related to Figure 2. **a**, Fraction of eggs on the lower sucrose option as a function of search duration with 95% confidence interval. **b**, Fraction of eggs on the lower sucrose option as a function of number of transitions during search with 95% confidence interval. **c**, Distribution of search durations. **d**, Distribution of number of transitions per search. In panels a-d, data was combined from 0/200, 0/500, and 200/500 mM chambers. 3979 eggs from 95 flies.

**Fig. S4.**
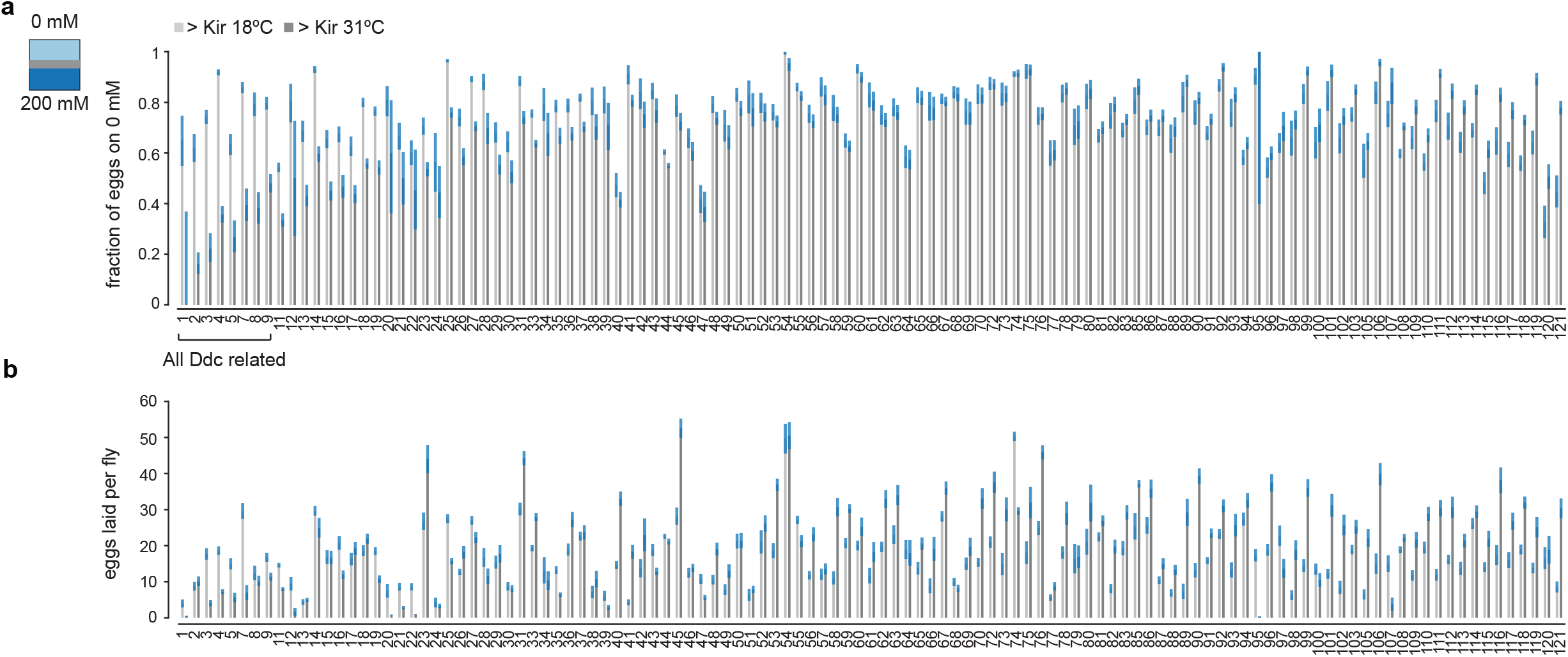
Summary of screen, related to Figure 6. **a**, Fraction of eggs on 0 mM with 95% confidence interval. Data is in increasing order of the x-axis in Fig. 6a. Genotype names are indicated in Table S2. **b**, Eggs laid per fly as in panel a.

**Fig. S5.**
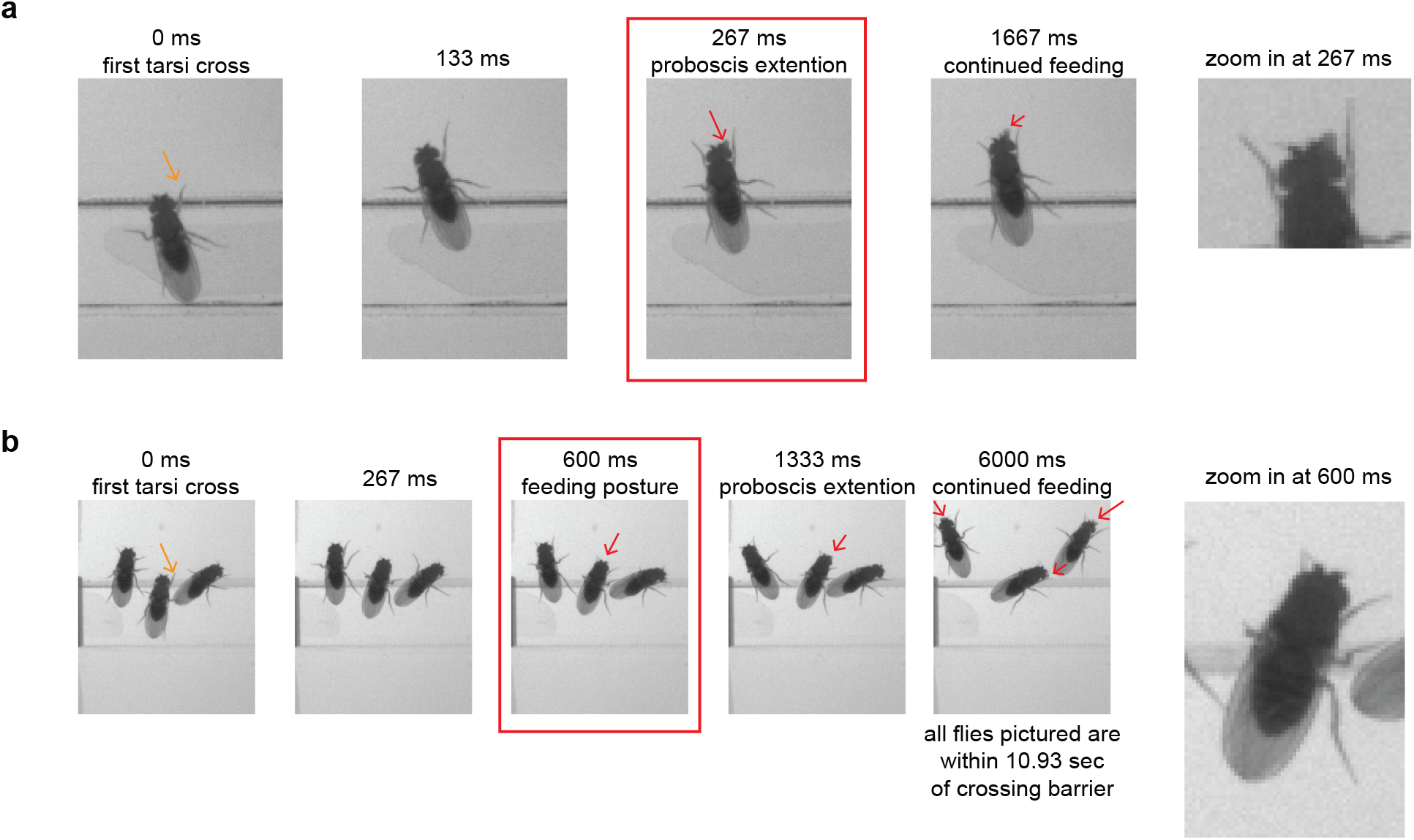
Example annotation of feeding events, related to Figure 6. **a**, Example annotation of a DDC > Kir (31°C pre) fly crossing the barrier from plain to 200 mM sucrose (orange arrow) and initiating proboscis extension (red square). The difference in time between the red square and orange arrow is the time used in Fig. 6b. **b**, Example annotation of a DDC > Kir (31°C pre) fly crossing the barrier from plain to 200 mM sucrose (orange arrow) and entering a feeding posture (red square). The difference in time between the red square and orange arrow is the time used in Fig. 6b. Proboscis extension events are invariably observed in subsequent frames once the proboscis is unobstructed by the head.

**Fig. S6.**
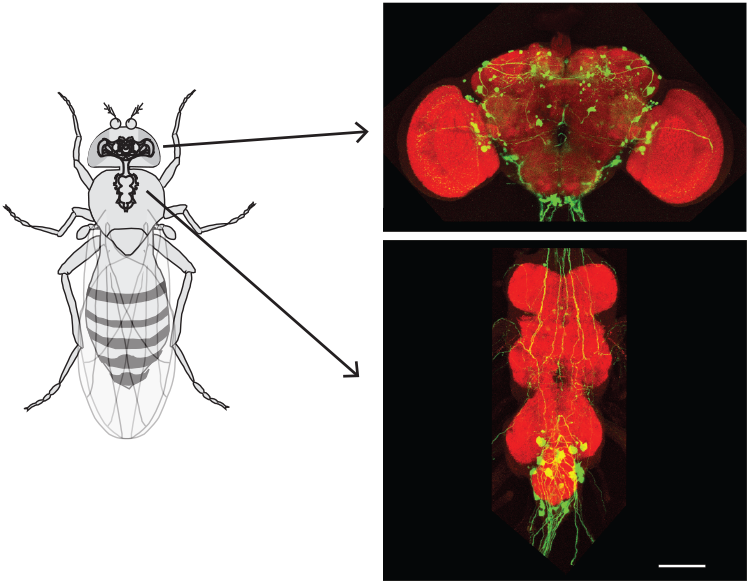
Neurons labeled by DDC-Gal4 in a female fly, related to Figure 6. Amplified EGFP signal (green) and nc82 counterstain (red). Fly genotype is w-; UAS 2xEGFP/+; DDC-Gal4/+. Scale bar, 100μm

### Supplemental Videos

**Video S1. Example egg laying event in a 0/500/200/500/200 mM sucrose (bottom to top) chamber at 10X speed, related to Figure 2**.

**Video S2. Example egg laying event in a 0/200 mM sucrose (bottom to top) 2 mm tunnel w/ bounce chamber at 10X speed, related to Figure 3**.

**Video S3. Example egg laying event in a 500/0/500 mM sucrose (bottom to top) chamber at 10X speed, related to Figure 4**.

### Supplemental Tables

**Table S1.** Summary of all tracking data, related to Figures 1 through 6.

| Genotype; Condition | Number of flies | Total number of eggs | Eggs laid on 0 mM | Eggs laid on 200 mM | Eggs laid on 500 mM | Total hours of tracking |
| --- | --- | --- | --- | --- | --- | --- |
| CS; 0 vs 200 | 47 | 1863 | 1711 | 152 | NA | 811 |
| CS; 0 vs 500 | 18 | 771 | 734 | NA | 37 | 304 |
| CS; 200 vs 500 | 30 | 1345 | NA | 1206 | 139 | 512 |
| CS; 0 vs 200 vs 200 | 24 | 1197 | 961 | 236 | NA | 406 |
| CS; 0 vs 200 vs 200 vs 200 vs 200 | 23 | 899 | 737 | 162 | NA | 377 |
| CS; 0 vs 500 vs 200 | 74 | 2313 | 1813 | 446 | 54 | 1270 |
| CS; 0 vs 500 vs 500 vs 500 vs 200 | 15 | 646 | 512 | 106 | 28 | 256 |
| CS; 0 vs 500 vs 200 vs 500 vs 200 | 21 | 878 | 632 | 223 | 23 | 334 |
| CS; 0 vs 200 circle chamber (4 mm) | 23 | 979 | 836 | 143 | NA | 416 |
| CS; 0 vs 200 circle chamber (2 mm) | 23 | 920 | 740 | 180 | NA | 412 |
| CS; 0 vs 200 circle chamber (2 mm w/ jump) | 17 | 675 | 413 | 262 | NA | 289 |
| CS; 500 vs 0 vs 500 | 24 | 943 | 758 | NA | 185 | 418 |
| CS; 0 vs 500 vs 500 mM | 18 | 749 | 721 | NA | 28 | 305 |
| CS; 500 vs 0 vs 200 | 63 | 1946 | 1127 | 668 | 151 | 1081 |
| DL; 0 vs 200 | 24 | 767 | 582 | 185 | NA | 403 |
| TUT; 0 vs 200 | 54 | 1185 | 723 | 462 | NA | 945 |
| D. suzukii; 0 vs 200 | 29 | 220 | 85 | 135 | NA | 507 |
| DDC > Kir 18 °C; 0 vs 200 | 48 | 1031 | 967 | 64 | NA | 834 |
| DDC > Kir 27 °C; 0 vs 200 | 20 | 762 | 700 | 62 | NA | 358 |
| DDC > Kir 29/29.5 °C; 0 vs 200 | 24 | 540 | 411 | 129 | NA | 427 |
| DDC > Kir 31 °C; 0 vs 200 | 41 | 533 | 158 | 375 | NA | 704 |

**Table S2.**
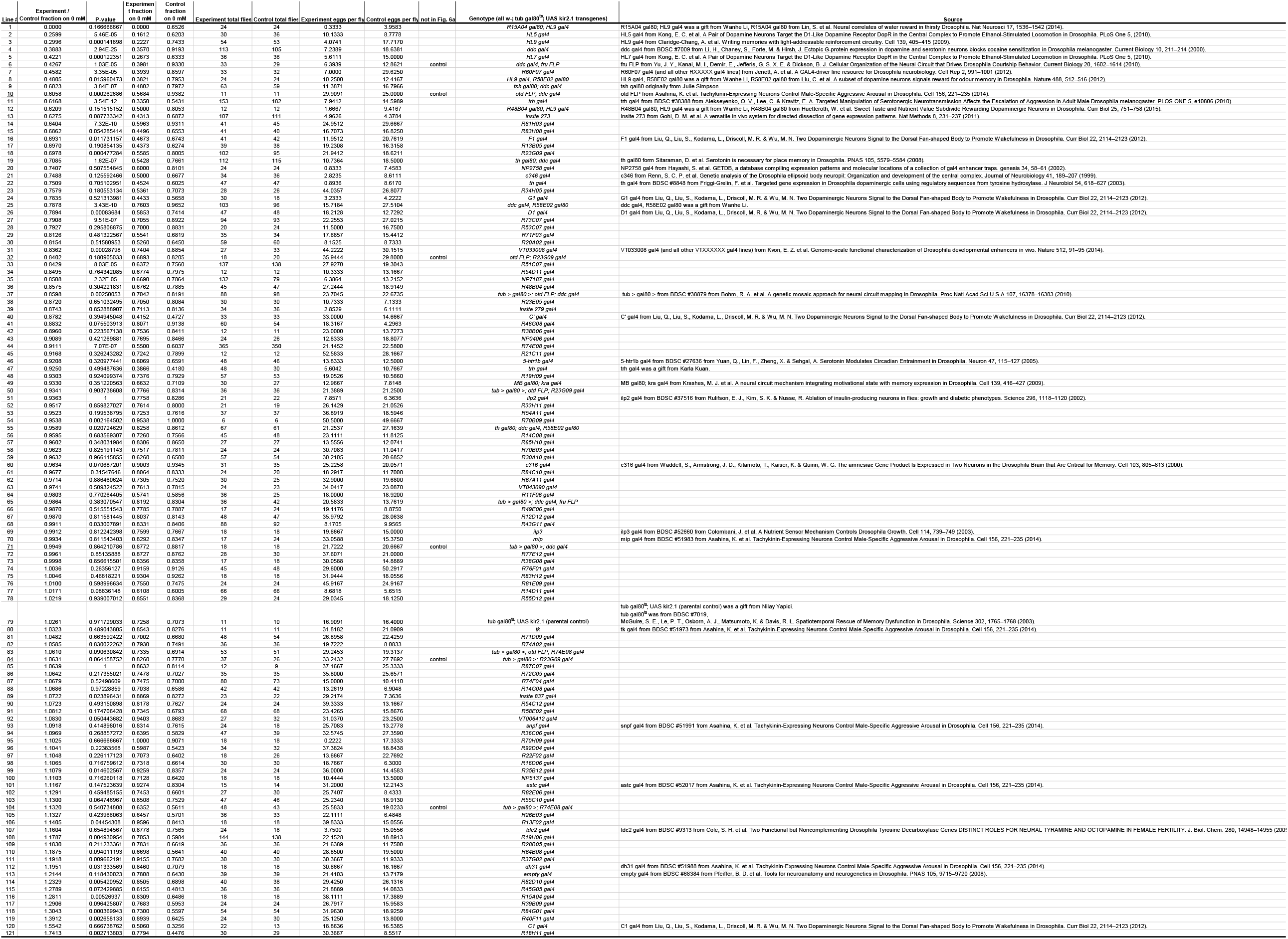
Summary of screen for egg-laying choice related neurons, related to Figure 6. All data in Fig. 6a is reported in increasing order of the x-axis in Fig. 6a. Data for 6 additional control lines that are not in Fig. 6a are also reported and marked ‘control’.

### Resource Availability

#### Lead Contact

Further information and requests for resources and reagents should be directed to the Lead Contact, Gaby Maimon.

#### Materials Availability

This study did not generate new unique reagents.

#### Data Availability

The datasets and code used in this study are available from the corresponding authors on request and will be made publicly available before or at time of publication.

### Experimental model and subject details

#### Flies

Flies were reared on a standard corn-meal medium at 25°C, ambient humidity, and a 12 h/12 h light/dark cycle. *D. suzukii* were reared with a wet Kimwipe (Kimberly Clark) in the vial or bottle with corn-meal medium. Canton S (CS) flies were obtained from Michael Dickinson and were originally from Martin Heisenberg. Dickinson Lab (DL) flies were originally from Michael Dickinson and were established in 1995 by interbreeding 200 iso-female wild caught stocks. Tuthill (TUT) flies were originally from John Tuthill and were established in 2009 by interbreeding 15 wild caught females^55^. *D. suzukii* flies were from Sylvia Durkin and Li Zhao and were originally the WT3 strain^56^. w-; tub Gal80^ts^; UAS Kir2.1 was a gift from Nilay Yapici. w-; UAS 2xEGFP was backcrossed into the Nippon-Project (NP) background (for a different study) and was originally from BDSC #6874^57^. The transgenic genotypes used for the screen, and their sources, are listed in Table S2.

### Method details

#### High-throughput egg-laying choice chamber

Experiments with wild type flies were conducted in the same manner as described in a previous study^13^. Experiments with Kir2.1 flies were conducted in the same manner to Kir2.1* assays from a previous study^13^, except flies were pre-treated at 18, 27, 29/29.5, or 31°C instead of only at 31°C. Dimensions and design of all chambers used in this study are shown in Fig. S1.

#### Gluing sensory organs

A thin layer of blue-light-cured glue (Bondic) was applied to sensory organs under gentle cold anesthesia. For the tarsi, only the distal two segments were lightly covered with glue. For the proboscis, the rostrum, proboscis, and labellum were glued such that the proboscis could not extend at all. Egg-laying assays were started 3 to 5 hours after glue application.

#### Latency to feeding assay

Flies were treated as in egg-laying assays except flies were mated in a bottle with only plain 0.8% agarose (a source of water, but not food). This treatment deprived flies of food for 26 hours allowing initiation of feeding to be measured as a proxy for the ability to sense sucrose. Groups of three to four females were placed on the 0 mM side of a 0 vs. 200 mM chamber under gentle cold anesthesia. These chambers were like the standard two choice egg-laying chambers except a sliding plastic barrier prevented flies from entering the 200 mM side of the chamber (Fig. S1h). Flies were imaged from above at 15 frames per second with a CM3-U3-13Y3M Chameleon camera (FLIR). A few minutes after starting image acquisition, we slid out the plastic barrier to allow flies to freely cross over to the 200 mM side of the chamber. The latency from the first time a tarsus crossed over the barrier to proboscis extension or entrance into a feeding posture (fly is still with head tilted down and forelegs forward)^58^ was scored manually (Fig. S5). All flies that were scored as entering a feeding posture were clearly also extending their proboscis once the proboscis became unobstructed by the head. Flies rarely, if ever, extended their proboscis or entered a feeding posture on 0 mM prior to crossing to 200 mM or when crossing from 0 mM to 0 mM in control chambers (data not shown).

#### Immunostaining and microscopy

The brain and ventral nerve cord of 2 to 5 day old females were dissected and processed similarly to a previous study^59^. Images were taken with a LSM 780 confocal microscope (Zeiss).

### Quantification and statistical analysis

#### Automated determination of egg-laying search

The egg-laying search was analyzed as described in a previous study^13^. We used an analysis of locomotor speed to define the start of the search (see below) and egg deposition to approximate the end of the search (which in free-walking flies in these chambers occurs only a few seconds after the search is concluded and the final abdomen bend to lay an egg is initiated^13^). The start of the search period was determined for each egg by smoothing the locomotor speed trace prior to egg deposition with an 18.5 s boxcar filter and identifying the first frame before an egg deposition event where this smoothed signal dropped below 0.1 mm/s. The minimum search duration was thus 18.5 s due to the length of the boxcar filter. These parameters were determined empirically to yield search onset times consistent with visual inspection of the data.

#### Calculation of egg-laying rate as a function of time

Egg-laying rate functions were calculated in the same manner as described previously^13^. Data from all flies tested in a given chamber type were combined prior to any calculations. First, we iterated through each time bin denoted on the x-axis and, for each bin, we counted the number of egg-deposition events that were assigned to that bin, #_eggs_(bin). Second, we iterated through the same time bins and counted the number of video frames in which flies were assigned to that bin, #_frames_(bin), during an egg-laying search period. Third, we iterated through the same time bins and counted the number of times flies changed assignment into that bin, #_visits_(bin), during an egg-laying search period (i.e., we did not keep incrementing the “visits” counter if the fly remained in a time bin from one frame to the next).

To get the mean egg-laying rate, we calculated #_eggs_/#_frames_ for each bin. Since videos were recorded at 2 fps, we multiplied the value for each bin by 120 to convert to units of eggs/minute.

To get the confidence interval per bin, we used the Clopper-Pearson method (‘exact’ binomial confidence interval) to calculate the 90% confidence interval for #_eggs_/#_visits_ for each bin. We then converted the confidence interval for each bin to units of eggs/minute by multiplying by 120*#_visits_/#_frames_ per bin. Note the confidence interval cannot be directly calculated from #_eggs_/#_frames_ because then the confidence interval would be dependent on the video framerate.

For these rate functions, search periods with duration less than 30 s were set to 30 s. This prevented very short search periods from introducing fluctuations in the rate curves (by contributing to the numerator and not contributing much to the denominator). As such, rate curves varied less from replicate-to-replicate or condition-to-condition. Note that search periods already had a minimum duration of 18.5 s as automatically generated by the search period calculation (Methods). Analyzing the data with no minimum search duration (or with other definitions of the search period) did not change any of the stated conclusions (Fig. S2). Binning the x-axis in different ways also did not qualitatively change any of our stated conclusions.

In this paper, we examine egg-laying rates as a function of time since visiting another substrate as our main approach for extracting algorithmic insight. As with any simple analysis, some important variables may be ignored (e.g., the locomotor vigor of the search bout, the duration of the visit to the previously visited substrate, visits to other substrates). These variables can vary in different chamber types and, as such, there are fluctuations in our egg-laying rate plots whose origin is not yet known. Nonetheless, this analysis allowed us to intuitively capture how the history of substrate experiences impacts the current egg-laying behavior.

#### Statistics

We used the two-sided Wilcoxon rank sum test to calculate all p-values. P-values for egg-laying rate curves or in the main text are comparing the number of trials with (or without) events in two separate groups. For a single group, trials with an event are treated as 1 and trials without an event are treated as 0. Then the two groups (each a set of 0 and 1) are compared using the two-sided Wilcoxon rank sum test (p-values calculated using the two-sided Fisher’s exact test are similar and similarly significant, except the 2^nd^ p-value in Fig. 3c is 5.8e-2). P-values in Fig. 3d are calculated by comparing the distributions in that figure. All other p-values are comparing two groups of individual flies, rather than individual trials.

Error bars are standard error of the mean unless otherwise described. For egg-laying choice fractions (like Fig. 1d or Fig. S3a) means are fraction of eggs laid on the lower sucrose option after all eggs from all flies are pooled and error bars indicate the 95% confidence interval of this fraction calculated using the Clopper-Pearson method (‘exact’ binomial confidence interval). For all experiments, no data were excluded, and no statistical method was used to choose sample size.

#### Data Analysis Software

All data analyses were done using MATLAB (MathWorks).

## Acknowledgements

We thank the Rockefeller University Precision Instrumentation Technologies facility for access to fabrication equipment; the Bloomington Drosophila Stock Center (NIH P400D018537), Li Zhao, Sylvia Durkin, Wanhe Li, Nilay Yapici, John Tuthill, Karla Kuan, and Mark Wu for fly stocks; Atsuko Adachi for help immunostaining many of the screened lines; and Nivedita Rangarajan, Sam Cohen, and Sylvia Durkin for assistance with experiments. Research reported in this publication was supported by a Brain Initiative grant from the National Institute of Neurological Disorders and Stroke (R01NS121904) to G.M. and Leon Levy Foundation fellowship and the Kavli Neural Systems Institute grant to V.V.. G.M. is a Howard Hughes Medical Institute Investigator.

## Author Contributions

V.V. and G.M. conceived the initial study and wrote the manuscript. V.V., with input from G.M., designed the experiments, performed the experiments, analyzed the data, interpreted the results, and decided on new experiments. Z.W., V.C., A.C., R.L., and S.L.S. assisted with performing various egg-laying experiments and interpreting the results. H.A. developed code to markedly speed up manual annotation of egg-deposition events.

## References

1. Karageorgi, M. et al. Evolution of Multiple Sensory Systems Drives Novel Egg-Laying Behavior in the Fruit Pest Drosophila suzukii. Current Biology 27, 847–853 (2017).

2. Zhu, E. Y., Guntur, A. R., He, R., Stern, U. & Yang, C.-H. Egg-laying demand induces aversion of UV light in Drosophila females. Curr Biol 24, 2797–2804 (2014).

3. Stensmyr, M. C. et al. A conserved dedicated olfactory circuit for detecting harmful microbes in Drosophila. Cell 151, 1345–1357 (2012).

4. Liu, W. et al. Enterococci Mediate the Oviposition Preference of Drosophila melanogaster through Sucrose Catabolism. Scientific Reports 7, 13420 (2017).

5. Yang, C., Belawat, P., Hafen, E., Jan, L. Y. & Jan, Y.-N. Drosophila Egg-Laying Site Selection as a System to Study Simple Decision-Making Processes. Science 319, 1679–1683 (2008).

6. Yang, C.-H., He, R. & Stern, U. Behavioral and Circuit Basis of Sucrose Rejection by Drosophila Females in a Simple Decision-Making Task. J. Neurosci. 35, 1396–1410 (2015).

7. Azanchi, R., Kaun, K. R. & Heberlein, U. Competing dopamine neurons drive oviposition choice for ethanol in Drosophila. Proc Natl Acad Sci U S A 110, 21153–21158 (2013).

8. Kacsoh, B. Z., Lynch, Z. R., Mortimer, N. T. & Schlenke, T. A. Fruit Flies Medicate Offspring After Seeing Parasites. Science 339, 947–950 (2013).

9. Joseph, R. M., Devineni, A. V., King, I. F. G. & Heberlein, U. Oviposition preference for and positional avoidance of acetic acid provide a model for competing behavioral drives in Drosophila. PNAS 106, 11352–11357 (2009).

10. Hussain, A. et al. Ionotropic Chemosensory Receptors Mediate the Taste and Smell of Polyamines. PLOS Biology 14, e1002454 (2016).

11. Dweck, H. K. M. et al. Olfactory Preference for Egg Laying on Citrus Substrates in Drosophila. Current Biology 23, 2472–2480 (2013).

12. Joseph, R. M. & Heberlein, U. Tissue-Specific Activation of a Single Gustatory Receptor Produces Opposing Behavioral Responses in Drosophila. Genetics 192, 521–532 (2012).

13. Vijayan, V. et al. A rise-to-threshold signal for a relative value deliberation. 2021.09.23.461548 https://www.biorxiv.org/content/10.1101/2021.09.23.461548v1 (2021) xdoi:10.1101/2021.09.23.461548.

14. Lin, C.-C., Prokop-Prigge, K. A., Preti, G. & Potter, C. J. Food odors trigger Drosophila males to deposit a pheromone that guides aggregation and female oviposition decisions. eLife 4, e08688 (2015).

15. Dennis, E. J., Goldman, O. V. & Vosshall, L. B. Aedes aegypti Mosquitoes Use Their Legs to Sense DEET on Contact. Curr Biol 29, 1551–1556.e5 (2019).

16. Wasserman, S., Salomon, A. & Frye, M. A. Drosophila Tracks Carbon Dioxide in Flight. Curr Biol 23, 10.1016/j.cub.2012.12.038 (2013).

17. Olsen, S. R. & Wilson, R. I. Lateral presynaptic inhibition mediates gain control in an olfactory circuit. Nature 452, 956–960 (2008).

18. Gou, B., Zhu, E., He, R., Stern, U. & Yang, C.-H. High Throughput Assay to Examine Egg-Laying Preferences of Individual Drosophila melanogaster. J Vis Exp (2016) doi:10.3791/53716.

19. Atallah, J., Teixeira, L., Salazar, R., Zaragoza, G. & Kopp, A. The making of a pest: the evolution of a fruit-penetrating ovipositor in Drosophila suzukii and related species. Proc Biol Sci 281, (2014).

20. Baines, R. A., Uhler, J. P., Thompson, A., Sweeney, S. T. & Bate, M. Altered Electrical Properties in Drosophila Neurons Developing without Synaptic Transmission. J. Neurosci. 21, 1523–1531 (2001).

21. McGuire, S. E., Le, P. T., Osborn, A. J., Matsumoto, K. & Davis, R. L. Spatiotemporal Rescue of Memory Dysfunction in Drosophila. Science 302, 1765–1768 (2003).

22. Wu, C.-L., Fu, T.-F., Chou, Y.-Y. & Yeh, S.-R. A Single Pair of Neurons Modulates Egg-Laying Decisions in Drosophila. PLOS ONE 10, e0121335 (2015).

23. Li, H., Chaney, S., Forte, M. & Hirsh, J. Ectopic G-protein expression in dopamine and serotonin neurons blocks cocaine sensitization in Drosophila melanogaster. Current Biology 10, 211–214 (2000).

24. Schultz, W., Dayan, P. & Montague, P. R. A Neural Substrate of Prediction and Reward. Science 275, 1593–1599 (1997).

25. Schultz, W. Getting formal with dopamine and reward. Neuron 36, 241–263 (2002).

26. Bayer, H. M. & Glimcher, P. W. Midbrain dopamine neurons encode a quantitative reward prediction error signal. Neuron 47, 129–141 (2005).

27. Chen, R. & Goldberg, J. H. Actor-critic reinforcement learning in the songbird. Current Opinion in Neurobiology 65, 1–9 (2020).

28. Montague, P. R., Dayan, P. & Sejnowski, T. J. A framework for mesencephalic dopamine systems based on predictive Hebbian learning. J Neurosci 16, 1936–1947 (1996).

29. Barto, A. G. Adaptive critics and the basal ganglia. in Models of information processing in the basal ganglia 215–232 (The MIT Press, 1995).

30. Sitaraman, D. et al. Serotonin is necessary for place memory in Drosophila. PNAS 105, 5579–5584 (2008).

31. Liu, C. et al. A subset of dopamine neurons signals reward for odour memory in Drosophila. Nature 488, 512–516 (2012).

32. Wang, F. et al. Neural circuitry linking mating and egg laying in Drosophila females. Nature 579, 101–105 (2020).

33. Webb, B. Cognition in insects. Philos Trans R Soc Lond B Biol Sci 367, 2715–2722 (2012).

34. Tolman, E. C. Cognitive maps in rats and men. Psychological Review 55, 189–208 (1948).

35. Tolman, E. C. & Honzik, C. H. Introduction and removal of reward, and maze performance in rats. University of California Publications in Psychology 4, 257–275 (1930).

36. Coen, P. et al. Dynamic sensory cues shape song structure in Drosophila. Nature 507, 233–237 (2014).

37. Gepner, R., Mihovilovic Skanata, M., Bernat, N. M., Kaplow, M. & Gershow, M. Computations underlying Drosophila photo-taxis, odor-taxis, and multi-sensory integration. eLife 4, e06229 (2015).

38. Neuser, K., Triphan, T., Mronz, M., Poeck, B. & Strauss, R. Analysis of a spatial orientation memory in Drosophila. Nature 453, 1244–1247 (2008).

39. Block, S. M., Segall, J. E. & Berg, H. C. Impulse responses in bacterial chemotaxis. Cell 31, 215–226 (1982).

40. Masson, J.-B., Voisinne, G., Wong-Ng, J., Celani, A. & Vergassola, M. Noninvasive inference of the molecular chemotactic response using bacterial trajectories. PNAS 109, 1802–1807 (2012).

41. Krakauer, J. W., Ghazanfar, A. A., Gomez-Marin, A., MacIver, M. A. & Poeppel, D. Neuroscience Needs Behavior: Correcting a Reductionist Bias. Neuron 93, 480–490 (2017).

42. Perugini, A., Ditterich, J. & Basso, M. A. Patients with Parkinson’s Disease Show Impaired Use of Priors in Conditions of Sensory Uncertainty. Curr Biol 26, 1902–1910 (2016).

43. Burke, C. J. et al. Layered reward signaling through octopamine and dopamine in Drosophila. Nature 492, 433–437 (2012).

44. Cohn, R., Morantte, I. & Ruta, V. Coordinated and Compartmentalized Neuromodulation Shapes Sensory Processing in Drosophila. Cell 163, 1742–1755 (2015).

45. Bennett, J. E. M., Philippides, A. & Nowotny, T. Learning with reinforcement prediction errors in a model of the Drosophila mushroom body. Nat Commun 12, 2569 (2021).

46. Berke, J. D. What does dopamine mean? Nat Neurosci 21, 787–793 (2018).

47. Eshel, N. et al. Arithmetic and local circuitry underlying dopamine prediction errors. Nature 525, 243–246 (2015).

48. Cohen, J. Y., Haesler, S., Vong, L., Lowell, B. B. & Uchida, N. Neuron-type-specific signals for reward and punishment in the ventral tegmental area. Nature 482, 85–88 (2012).

49. Watabe-Uchida, M., Eshel, N. & Uchida, N. Neural Circuitry of Reward Prediction Error. Annu Rev Neurosci 40, 373–394 (2017).

50. Haber, S. N. The place of dopamine in the cortico-basal ganglia circuit. Neuroscience 282, 248–257 (2014).

51. Dayan, P. & Niv, Y. Reinforcement learning: The Good, The Bad and The Ugly. Current Opinion in Neurobiology 18, 185–196 (2008).

52. Scheffer, L. K. et al. A connectome and analysis of the adult Drosophila central brain. eLife 9, e57443 (2020).

53. Zheng, Z. et al. A Complete Electron Microscopy Volume of the Brain of Adult Drosophila melanogaster. Cell 174, 730–743.e22 (2018).

54. Venken, K. J. T., Simpson, J. H. & Bellen, H. J. Genetic manipulation of genes and cells in the nervous system of the fruit fly. Neuron 72, 202–230 (2011).

55. Tuthill, J. C., Chiappe, M. E. & Reiser, M. B. Neural correlates of illusory motion perception in Drosophila. PNAS 108, 9685–9690 (2011).

56. Chiu, J. C. et al. Genome of Drosophila suzukii, the spotted wing drosophila. G3 (Bethesda) 3, 2257–2271 (2013).

57. Halfon, M. S. et al. New fluorescent protein reporters for use with the drosophila gal4 expression system and for vital detection of balancer chromosomes. genesis 34, 135–138 (2002).

58. Flood, T. F. et al. A single pair of interneurons commands the Drosophila feeding motor program. Nature 499, 83–87 (2013).

59. Jenett, A. et al. A GAL4-driver line resource for Drosophila neurobiology. Cell Rep 2, 991–1001 (2012).

